# Intrinsic Population Dynamics are a Neuronal Substrate for Visual Attention

**DOI:** 10.64898/2026.06.02.729565

**Authors:** Florian H. Schmidt, Wiktor Młynarski, Abir George, Anton Sumser, Gašper Tkačik, Maximilian Joesch

## Abstract

Perception results from a dynamic interplay between the feedforward processing of sensory stimuli and intrinsic neural activity ^1^, which is often dismissed as noise ^2^. To tailor perceptual processes to the organism’s current needs on a continuous, moment-to-moment basis, intrinsic dynamics – rather than just being noise – have been suggested to reflect prior expectations ^3,4^, task demands ^5^, and attentional focus ^6^. Here, we identify a novel signature of attentive state in which intrinsic, collective neural activity modulates visuospatial attention by dynamically interacting with sensory input within the superior colliculus (SC), a midbrain hub that integrates bottom-up visual input with top-down signals ^7^. We show that these intrinsic dynamics organize cell activity into structured, blob-like features that are temporally and topographically localized and that rival sensory-evoked responses in strength. These features emerge as animals learn to engage in the visual detection task, and predict behavioral outcomes on a trial-by-trial basis. Although independent of sensory input or overt behavior, the features can be recruited to enhance visual responses four-fold in the attentive state, generating a “dynamic saliency map” that aligns with reaction times and behavioral outcomes. A computational model indicates that tunable blob-like features can arise from local excitatory–inhibitory interactions within the SC, and enhance sensory-evoked responses in line with observations. Together, our results identify state-dependent intrinsic population activity that interacts with sensory input to give rise to a saliency-map-like representation linked to flexible, goal-directed behavior.

## Main

In the absence of external sensory stimuli, the brain is constantly active and consumes most of its energy ^8^. This spontaneous neural activity has often been considered to be noise — random and irregular fluctuations that obscure and interfere with the sensory signal ^2,9^. Thus, to differentiate the “true signal” from “noise”, sensory physiology typically focuses on the average response, assessing reliability through the signal-to-noise ratio. However, in an ever-changing environment, perception cannot depend on trial-averaged information; instead, it must rely on moment-to-moment assessments of the environment. These assessments are strongly influenced by the current internal states, context, and task demands. An alternative explanation for “noise” is therefore simply our ignorance about such internal processes: in fact, they may support rather than distort the transmission and processing of relevant information ^10,11^.

Experimental results, primarily in cortical structures, provide evidence for this view ^3–5,11–13^. For example, spontaneous neuronal dynamics have been shown to be structured in time and space ^1,3,14,15^, reflect behavior ^5^ and show systematic temporal dependencies ^16^. Locally synchronous membrane depolarizations regulate the amplitude and spread of sensory-evoked signals ^17^ and spontaneous cortical traveling waves influence perception ^1^. In addition, visual processing has been shown to activate neuronal ensembles that may also coactivate spontaneously due to the underlying circuit connectivity ^4,13,18^, a property that is thought to represent prior expectations about the environment ^3^.

Among the many internal processes that contribute to variability, selective visual attention is particularly prominent ^10^. Selective visual attention—the prioritization of behaviorally relevant regions of stimulus space for enhanced processing—is a fundamental cognitive function^19,20^ dynamically shaped by current task demands. The best neuronal correlate of this function has been found in the enhanced response of visual cortical neurons to attended stimuli ^20^. While cortical pathways have been extensively studied, subcortical pathways via the superior colliculus (SC) also contribute to visual attention through mechanisms that are distinct from the classical cortical effects ^7,21^. The SC is particularly well positioned to implement such sensory selection. It integrates bottom-up retinal input with contextual and top-down signals, both directly from cortical areas^22^ and through neuromodulatory centers^23–27^, while simultaneously influencing orienting and goal-directed behaviors^28^. Consistent with this central role, the SC has been implicated in action generation, decision-making, and attentional control ^7,29^. In primates, these interactions are thought to give rise to a saliency map — a localized pattern of neuronal activity that guides the gaze toward behaviorally relevant targets in the visual scene ^28,30^.

Here, we identify a previously unrecognized form of intrinsic collective neuronal activity in the mouse superficial SC (sSC). We establish a direct link between this activity and an attentive, task-engaged behavioral state that dynamically modulates visuospatial processing. These collective dynamics are spatially and temporally structured and emerge in a state- and attention-dependent manner, for example through the association of spatial visual stimuli with reward. Although they are stochastic and independent of overt behavior or reward, they locally enhance behaviorally relevant visual responses. Notably, these dynamics predict attentive states that emerge with learning of the association between visual space and reward, and track behavioral performance and reaction times. They further interact with visually driven activity to generate what we term a “dynamic saliency map”, which underpins goal-directed actions. Computational modeling suggests that such collective activity can arise from intrinsic excitatory–inhibitory interactions within the SC, rendering it sensitive to internal attentive state changes, e.g., those driven by neuromodulatory influences ^31^. Importantly, an emergent property of these interactions is the sensory-driven recruitment of intrinsic activity observed experimentally. Together, our findings uncover a collective dynamical brain state essential for continuously prioritizing and acting upon behaviorally relevant features of the environment.

## Results

### Visual spatial attention enhances sensory representations in the SC

Because the SC lies at the interface between visual processing, internal-state modulation, and orienting behavior, we asked whether population activity in the SC can be related to attentive behavior. Inspired by previous work ^32–34^, we first established a spatial visual detection task that elicits behavioral signatures consistent with spatial attention effects and is also fast to learn, allowing for longitudinal imaging experiments (Fig. 1a,b). Mice (n = 17) were trained to detect a faint visual stimulus presented at different spatial locations and to report its detection by licking. The first lick during the reward window triggered reward delivery after ∼350 ms. Non-task-related licking was discouraged with a timeout and mild auditory penalty (Fig. 1b), triggered when licking exceeded 0.2 licks s⁻¹ before stimulus onset. Animals learned to suppress spontaneous licking and respond reliably within ∼10 sessions (Fig. 1c,d), while remaining engaged for at least 120 trials per session (Suppl. Fig. 1a).

**Fig. 1.**
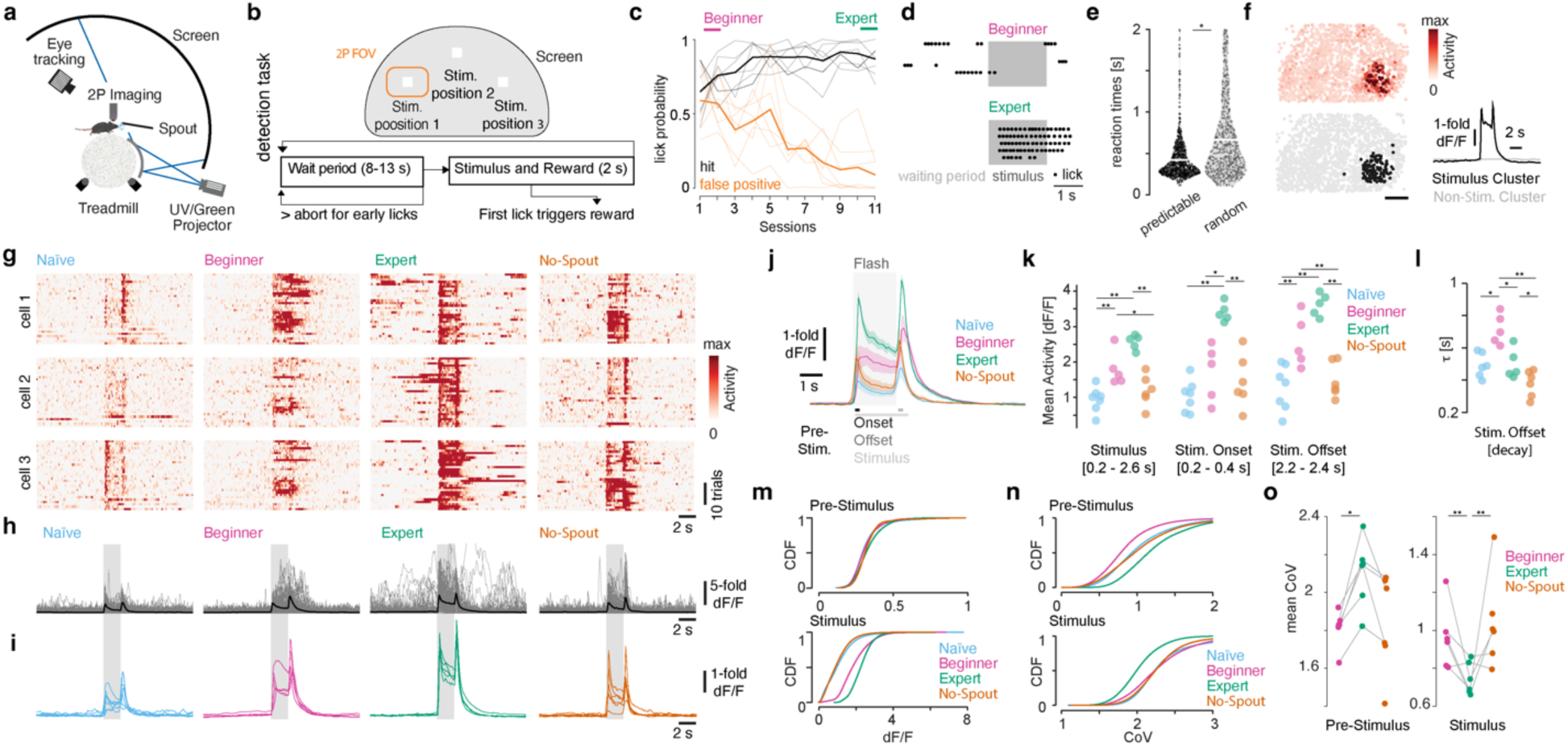
Visual responses in the sSC change across learning stages and internal state during a visual attention task. **a.** Schematic of the two-photon imaging and behavioral training setup. Mice are head-restrained while running on a spherical treadmill, with dim visual stimuli presented on a panoramic dome. **b.** Top: Schematic of the panoramic dome setup showing three different stimulus locations and the corresponding retinotopic area imaged in the superior colliculus (SC). Bottom: Visual spatial attention detection task. Rewards for detection are delivered if the animal licks during the stimulus presentation, which is triggered by the first lick. **c.** Licking behavior across learning stages. Thin lines indicate individual animals; thick lines represent the mean. “Beginner” and “Expert” stages are depicted. **d.** Example of licking behavior across learning stages for a representative animal. **e.** Reaction times for trials with stimuli presented either at a fixed location (“predictable”) or randomized across locations (“random”). **f.** Example imaging field-of-view from the SC showing the average response to a visual flash (top), with k-means clustering distinguishing visually responsive (bottom black) and non-responsive (not stimulated) cells (bottom grey). Right: Temporal response profiles of each cluster. **g.** Calcium activity traces from representative visually responsive cells across learning stages. **h.** Example trials from (**g**) (gray) and their mean (black) illustrate increased intrinsic activity across learning stages. **i.** Average responses of all visually responsive cells per animal and learning stage. **j.** Population average responses across animals and learning stages. **k.** Quantification of mean responses per animal, corresponding to panel (**i**). **l.** Quantification of the decay time constant from traces shown in (**i**). **m.** Cumulative distribution of neural activity across learning stages. **n.** Same as (**m**), but showing the activity coefficient of variation (CoV). **o.** Relative change in mean CoV compared to the beginner stage for each animal. All statistical details are provided in Suppl. Table 1.

As expected from previous studies ^32,33^, behavioral performance revealed hallmarks of visual spatial attention: reaction times were significantly faster when the stimulus was presented repeatedly at the same location (predictable condition) compared to randomized stimulus locations (Fig. 1e). Conversely, reaction times were markedly slower when the stimulus location changed after repeated presentations (“switch trials”), but became faster when a weak global cue was provided (Suppl. Fig. 1b-d). These results were independent of arousal, as measured by pupil dynamics (Suppl. Fig. 1e-g). Moreover, decreasing signal strength by reducing stimulus size led to a corresponding increase in reaction times and a decrease in hit rates (Suppl. Fig. 1h-j). We observed two forms of licking: isolated licks and short bursts, defined as consecutive licks separated by less than 200 ms. Although the first lick during the stimulus period triggered reward delivery, bursts of at least four licks typically preceded reward arrival, consistent with expectation of the go-cue–associated outcome (Suppl. Fig. 1k). Bursts were rare during pre-stimulus periods in trained animals, and early-session timeouts were primarily caused by accumulated isolated licks, indicating stable false-alarm rates, impulsivity, and decision criteria. Together, these results show that the paradigm rapidly trains mice to flexibly allocate attentional resources across visual space according to task demands within a timescale compatible with longitudinal imaging.

To examine visual responses during training progression, we expressed the calcium indicator GCaMP6f^35^ transgenically in a Cre-dependent manner using the Rorb-Cre line, which labels a diverse population of neurons in the sSC ^36,37^. This approach ensured consistent expression levels across imaging sessions and enabled us to monitor 2260 ± 1146 neurons per animal (mean ± STD across n = 8 mice). The animals were head-fixed on a spherical treadmill in front of a custom-built spherical dome projection system (Fig. 1a). Note that only one of the several stimuli fell within the imaged field of view (FOV) (Fig. 1b). We classified visually responsive and non-responsive (not stimulated) neurons using k-means clustering (Fig. 1f). As expected, responsive neurons exhibited average responses to both stimulus onset and offset (Fig. 1f, bottom right). We used this clustering method to analyze visual responses across distinct learning stages and behavioral conditions of the predictable condition: naïve (before training), beginner (first training session), expert (after suppression of non-task-related licking), and no-spout (expert animals prevented from engaging in the task by removing the reward spout) (Fig. 1e,g,h). These experiments revealed a pronounced increase in visual response strength across learning stages, at the single-cell (Fig. 1g) and population levels (Fig. 1h), as well as across animals (Fig. 1i), with responses approximately doubling on average from naïve to beginner, and again from beginner to expert. Importantly, response strength receded to the naïve stage in the no-spout condition, indicating that these changes arise due to task-relevant states rather than representational changes driven by learning (Fig. 1j,k). They were furthermore not caused by task-irrelevant movements, reward or licks (Suppl. Fig. 2, Supplementary Video 1). We therefore operationally defined “attentive states” based on these observable changes, so that naïve animals exhibited low attentional engagement, beginners showed a transitional increase, and experts maintained a stable, high-attentive state that dissipated when task engagement was prevented in the no-spout condition.

As expected from a feed-forward sensory system, the average response was a good descriptor of single trial responses in naïve animals. However, this pattern markedly changed with learning progression. In beginners, activity increased after stimulus offset, resulting in a prolonged decay of the average offset response (Fig. 1l). In experts, similarly pronounced increases occurred at random times during the trial, presenting as seemingly uncorrelated noise across the session (Fig. 1g,h). As a result, these random increases were not apparent in the session average (Fig. 1i) or in the distribution of prestimulus activity (Fig. 1m, top), but became evident in the increased prestimulus coefficient of variation, a measure of response dispersion (Fig. 1n, top). On the contrary, during visual stimulation in experts, stimulated cells showed both enhanced activity and a decrease of coefficient of variation in aggregate (Fig. 1m,n, bottom).

A more detailed, animal-by-animal analysis confirmed that expert visual representations became more stable compared to the beginner stage and the no-spout condition (Fig. 1o).

### Intrinsic dynamics are collective and change with attentive states

The prominence and magnitude of intrinsic, non-sensory activity (“noise”) were comparable to visually evoked responses (“signal”), rendering them nearly indistinguishable without knowledge of stimulus timing, particularly in the expert stage (Fig. 1g,h; see expert). We therefore sought to characterize the structure of this intrinsic activity beyond the visually responsive cell cluster (Fig. 1f). To do so, we used population entropy (PE) - a measure that is maximal when activity is broadly distributed across the neuronal population and minimal when concentrated in a single neuron (see Methods). PE thus captures changes in how population activity is distributed across neurons (Fig. 2a,b), providing information complementary to measures of mean or total activity. While independent of the spatial arrangement of active cells, this metric detects moment-to-moment changes in collective dynamics on a single-trial basis. On the one hand, PE underwent a pronounced decrease upon stimulus presentation (Suppl. Fig. 3a,b). As expected based on averaged response, this decrease was accompanied by concurrent spatial clustering of active cells (Suppl. Fig. 3c,d). On the other hand, decreases in PE also enabled us to uncover previously unrecognized, single-trial dynamical features in non-sensory activity, which we term “blobs” (Fig. 2a–e, top, Suppl. Fig. 3e). Blobs are large-amplitude (Fig. 2c,e), spatially localized neural activity patterns (Fig. 2g), distributed across the visual space (Suppl. Fig. 3e,f, Supplementary Video 1), and appearing consistently across animals (Fig. 2h). The appearance of such localized intrinsic activity patterns is particularly notable in the SC, whose retinotopic organization and established role in stimulus selection suggest that local population dynamics may have direct consequences for the prioritization of sensory information.

**Fig. 2.**
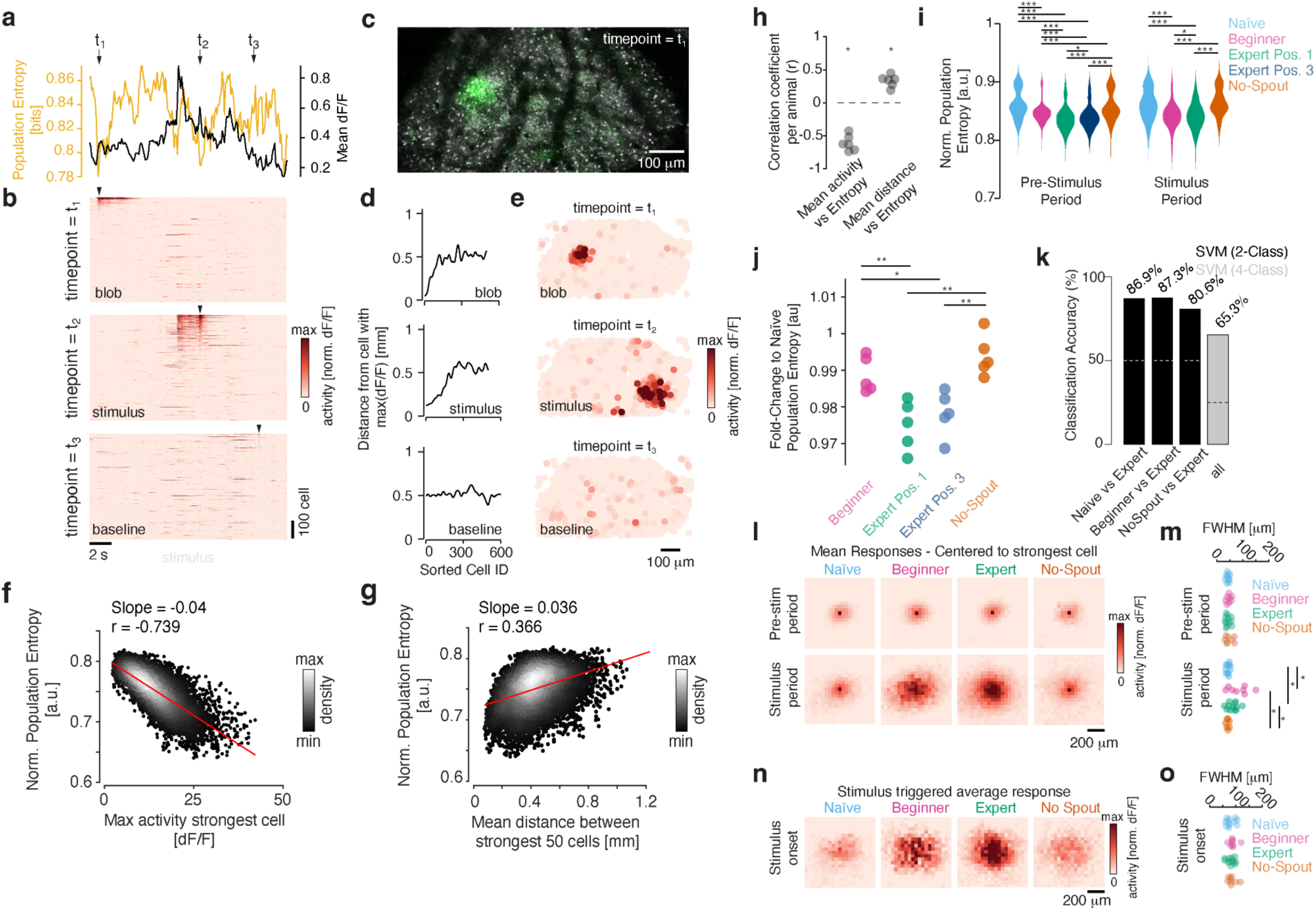
Spatially-structured intrinsic sSC dynamics dominate single-trial activity of engaged expert animals. **a.** Representative session showing single-trial population entropy (PE, yellow) and mean calcium activity across all cells (black) in an expert animal. **b.** Activity from the session in (**a**), sorted by strength for three selected timepoints (*t₁, t2, t3*), as indicated by the arrowheads. **c.** Maximum intensity projection (grey) superimposed with the raw Calcium activity (green) of unprocessed imaging data at timepoint *t1*. **d.** Smoothed spatial distance between the maximally active cell and cells ordered by activity, sorted as in (**b**), highlighting the spatial organization of activity patterns. **e.** Spatial map of activity for each cell at timepoint *t*, revealing an activity blob at t1 and the visually responsive cell cluster at t2. **f-g.** Scatter plots and linear fits showing the relationship between PE and the maximally responsive cell (**f**) and the average distance of the 50 strongest cells (**g**). 50 cells were chosen because they represent the lower bound on the number of neurons typically participating in a blob, see (**b**) (at *t1*). **h.** Correlation coefficients, as in (**f-g**), calculated for individual animals. **i.** Distribution of PE across conditions. **j.** Fold-changes in PE relative to Naïve, per animal. **k.** 5-fold cross-validated classification accuracy of a kernelized SVM trained to distinguish learning stages based on a time window of PE dynamics, using two- and four-class classifiers. **l.** Top: Average rasterized spatial activity (40 x 40 x 250 µm voxels), centered on the maximally active voxel, for each low-PE event (as in panel b, timepoint *t₁*). Bottom: Same analysis as in the top panel, but for the stimulus onset timepoint *t2*. **m.** Full-width at half-maximum (FWHM) of the activity patterns shown in (**l**), calculated per animal and condition. **n.** Average spatial activity, centered on the peak responding voxel. **o.** FWHM of the visual response shown in (**n**), calculated per animal and learning stage. All statistical details are provided in Suppl. Table 1.

The PE distribution changed systematically across learning stages, driven by a marked increase in the prevalence of blob-associated low-PE events from the naïve to the expert stage, independent of stimulus position. The distribution reverted to naïve-like shape under the no-spout condition (Fig. 2i,j; Supplementary Video 1). Its independence from stimulus laterality suggested that the observed dynamics were not contingent on direct SC activation. While low-PE events were not correlated with any overt behavioral parameter (Suppl. Fig. 4), already a 2 s window of single-trial PE dynamics preceding stimulus onset served as an excellent predictor of learning stage, as demonstrated by a support vector machine (SVM) classifier (Fig. 2k).

We next took advantage of the fact that low-PE events served as indicators of coordinated intrinsic activity whose spatial properties we wished to characterize. By averaging single-trial spatial activity maps centered on the maximally active cell at low-PE events, we identified blob-like activity patterns with remarkably similar spatial properties across learning stages (Fig. 2l,m, top). This suggested that learning primarily alters the frequency of blob occurrence rather than their spatial structure.

We then repeated the same analysis during the stimulus period (Fig. 2l,m, bottom). In the naïve stage and under the no-spout condition, the resulting spatial activity closely resembled the pre-stimulus blobs, suggesting that visually evoked responses are comparable in magnitude to intrinsic collective activity patterns. In contrast, during the beginner and expert stages, low-PE-triggered spatial activity was markedly enhanced, suggesting that sensory input selectively recruits intrinsic events. Consistent with this interpretation, stimulus-triggered average responses resembled those identified in our previous analyses, exhibiting similar mean but more variable full-width at half-maximum (FWHM) values (Fig. 2n,o). Together, these observations suggest that intrinsic blob-like activity locally co-occurs with, and thereby amplifies, sensory-evoked responses. More broadly, the observations establish intrinsic dynamics as a defining feature of the animal’s attentive state.

### Intrinsic dynamics are organized into blob-like features recruited by the stimulus

While the intrinsic dynamics define the animal’s attentive state, it remains unclear whether the emergence of blob-like features plays a predictive role, in which blobs act as spatially localized attentional “spotlights” that selectively bias future sensory processing. To address this question, we decomposed neuronal activity into spatiotemporal motifs using non-negative matrix factorization (NMF) (Fig. 3a, Suppl. Fig. 5a–f; see Methods). This approach enabled us to isolate and track individual motifs within the ongoing activity at a single-trial level. Given the strong correlations among nearby neurons (Suppl. Fig. 5d), we discretized the imaged region into a regular lattice of 13 × 26 voxels (40 × 40 × 250 μm³ each), including only voxels containing more than five neurons (Suppl. Fig. 5c, Supplementary Video 2). Activity was averaged within each voxel, and NMF decomposition of this voxelized signal accounted for ∼50% of the total variance, with comparable single-trial reconstruction errors across learning stages (Suppl. Fig. 5g,h). Reconstructions closely matched the recorded data (Suppl. Fig. 5f,i–l, Supplementary Video 3) and revealed spatially localized components underlying intrinsic dynamics that strongly resembled those identified using the population entropy metric (Fig. 2l, Suppl. Fig. 5m,n). The average temporal profile of these intrinsic features showed that blobs appear, persist, and disappear with identical kinetics across learning stages, with comparable rise and decay times of 2.85 ± 0.10 s (mean ± SEM, n = 39 recordings across conditions; Suppl. Fig. 5o, p). Overall, these results validated our previous findings based on PE (Fig. 2), including their independence of licks, reward, or other behavioral parameters (Suppl. Fig. 6). Consistently, low-PE events closely aligned with peaks in the NMF-derived temporal components, lagging by only ∼100 ms on average (median; Suppl. Fig. 5q).

**Fig. 3.**
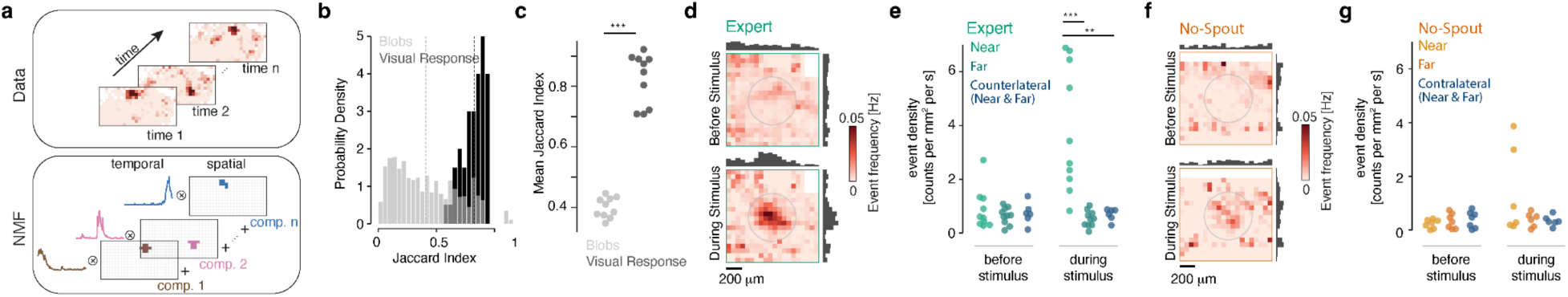
Uniformly distributed blob-like features become localized and strongly amplified during visual stimulation in the attentive state. **a.** Schematic of NMF decomposition. The data are decomposed into spatiotemporal components whose sum reconstructs the original activity. **b.** Distribution of Jaccard indices between cell assemblies responding to the stimulus (black) and those participating in NMF-derived blob-like features (gray). **c.** Mean Jaccard index from (**b**) across animals. **d.** Probability map of blob location (defined as their center of mass) before and during visual stimulation in expert animals, with corresponding marginal distributions. **e.** Area-normalized event occurrence frequency near (inside the faint grey circle in (**d**)), far (outside), or across the entire field during contralateral stimulation. **f–g**. Same as (**d, e**), but for the no-spout condition. All statistical details are provided in Suppl. Table 1.

Leveraging the temporal and spatial resolution provided by the NMF decomposition (Fig. 3a), we next asked whether the blob-like features reflect a fixed modular tiling of the SC ^38^ or instead arise from flexible population assemblies in which individual neurons participate in multiple distinct blobs. To address this, we computed the Jaccard index, a measure of set overlap that quantifies the similarity in neuronal participation across blobs (see Methods). Compared to visually evoked responses, intrinsic blobs exhibited markedly lower Jaccard indices, indicating that neurons were flexibly recruited across different events (Fig. 3b). Mean Jaccard indices were highly consistent across animals and clearly distinct from those associated with retinotopically evoked visual responses (Fig. 3c).

We next asked whether blobs could act as a spatial “spotlight,” biasing their probability of occurrence toward specific SC locations prior to stimulus onset, e.g., at the location of the expected stimulus. To isolate intrinsic dynamics while attenuating stimulus-locked components, we subtracted the mean visually evoked response from individual trials. Because intrinsic activity manifested as trial-specific, spatially-variable events superimposed on sensory responses, this procedure largely preserved intrinsic dynamics while reducing the average evoked component. We then defined the center of mass of each blob and computed spatial occurrence maps relative to the stimulus location (Fig. 3d–g). Before stimulus onset, blob occurrences were evenly distributed across the SC surface in both the expert stage and the no-spout condition, indicating that intrinsic dynamics did not spatially bias blob location. However, they did modulate overall occurrence probability, with fewer blobs observed in the no-spout condition, consistent with earlier findings (Fig. 2). In contrast, during stimulus presentation, blob occurrence increased markedly at the stimulus location in expert animals (Fig. 3d, bottom; e), an effect that was strongly attenuated in the no-spout condition (Fig. 3f, bottom; g). These changes were not explained by reward or behavior, as rewarded stimuli presented on the opposite side did not alter blob occurrence in the imaged hemisphere (Fig. 3e,g). The enhanced visual responses (Fig. 1) are therefore likely mediated by the emergence of blobs at the stimulus location, indicating that for expert animals in the attentive state behaviorally relevant visual stimuli can selectively recruit local intrinsic dynamics.

### Intrinsic dynamics predict performance and reaction times in visual spatial attention task

To test whether blob recruitment contributes to behavioral performance, we compared successful trials (hits) with trials in which animals licked after the reward window or failed to lick altogether (misses). This analysis revealed a pronounced enhancement of visual responses during hit trials that closely tracked behavioral performance at the single-cell (Fig. 4a), population (Fig. 4b,c), and across-animal levels (Fig. 4d). On average, visual responses were 34.9% weaker during miss compared to hit trials. Interestingly, the main differences between hit and miss trials were apparent not only immediately after stimulus onset but also approximately one second later, suggesting a non-sensory component (Fig. 4b). These effects were not attributable to behavioral differences, reward delivery, or licking (Suppl. Fig. 6; Supplementary Video 3 & 4). Interestingly, hit and miss trials could be predicted prior to visual stimulation, similarly to how learning stages could be predicted (Fig. 2k), with a SVM classifier (Fig. 4e). While the classifier showed limited accuracy in distinguishing closely matched hit and miss trials, classification performance improved markedly for miss trials with longer reaction times, indicating that pre-stimulus internal state is predictive of behavioral outcome.

**Fig. 4.**
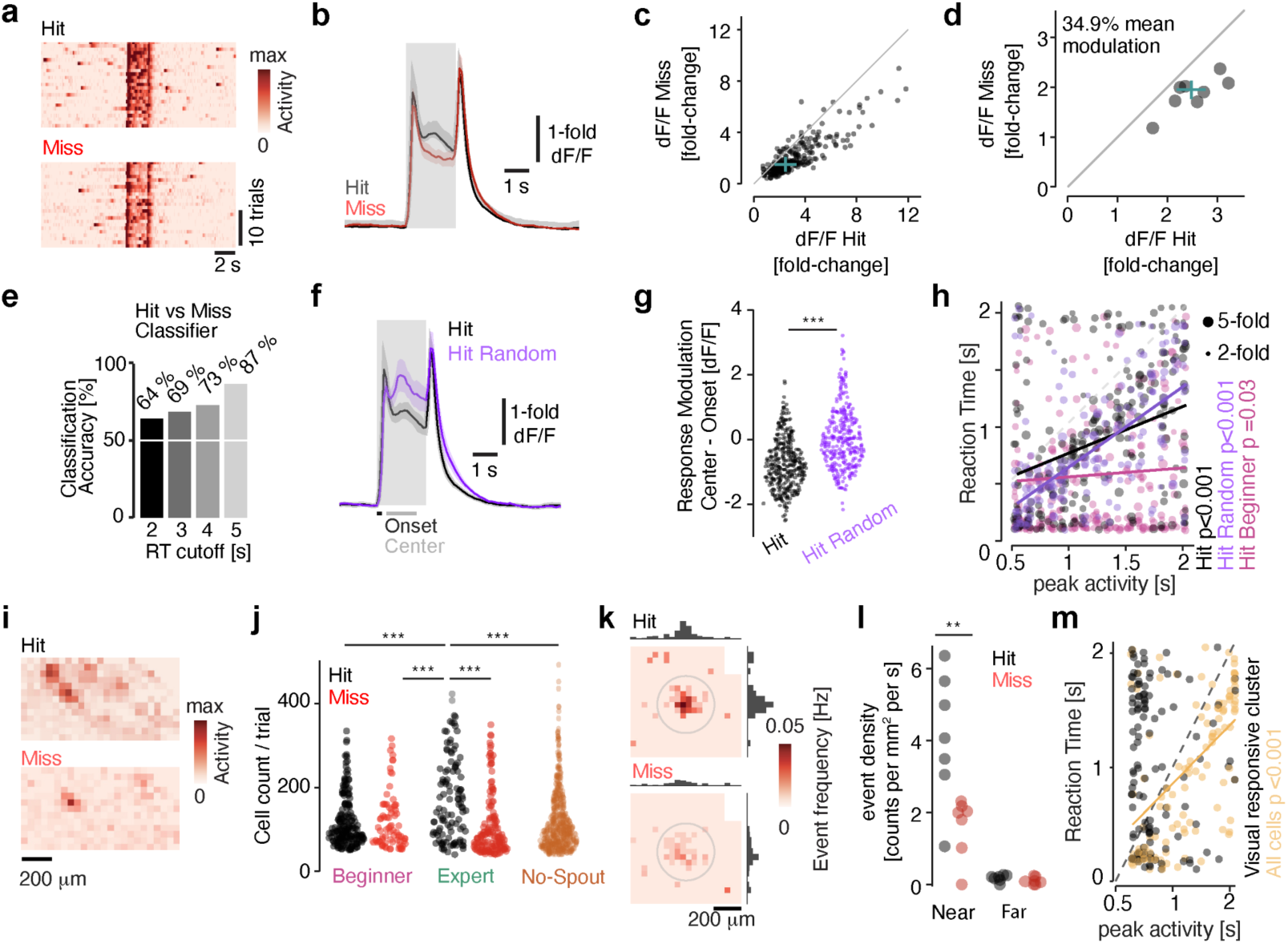
Hit and miss trials link intrinsic dynamics with behavioral performance. **a-b.** Representative calcium traces from a visually responsive cell (**a**) and population average (**b**) for predictable hit and miss trials. **c-d**. Mean visual responses of visually responsive cells for one expert animal (**c**) and across animals (**d**); crosses indicate median ± MAD. **e.** SVM classification of hit versus miss trials using a 2 s pre-stimulus window of single-trial population entropy; late misses (longer RTs) showed improved classification. **f.** Population-averaged responses for predictable and random trial setups on hit and miss trials. **g.** Modulation index of visual responses during hit trials under predictable and random experimental conditions. **h.** Relationship between peak response timing of visually responsive clusters and reaction times for predictable (expert and beginner) and random (expert) conditions. Dot size indicates response magnitude; solid lines show significant linear fits. **i.** Single-trial activity maps during presentation of a small (2° visual field) stimulus for representative hit and miss trials. **j.** Quantification of stimulus-driven cluster sizes across learning stages for hit and miss trials. **k.** Occurrence frequency maps of blob centers-of-mass during visual stimulation in expert animals for hit and miss trials, with marginal distributions. **l.** Area-normalized event density near versus far from the stimulus location. **m.** Relationship between peak activity timing of visually responsive clusters (or all cells, respectively) with reaction times during small-stimulus experiments. Solid line shows a significant linear fit. All statistical details are provided in Supplementary Table 1.

We next asked whether changing the predictability of sensory input, and thus the attentional demands of the task, modulated the intrinsic activity patterns. To test this, we varied stimulus location from predictable to random, thereby increasing spatial uncertainty and prolonging reaction times (Fig. 1e). Interestingly, in hit trials during random stimulus presentation (“hit random”), the intrinsic activity was stronger overall, and tended to occur later during the stimulus presentation (Fig. 4f), a change that was highly significant (Fig. 4g). To determine whether such intrinsic activity was instructive, we correlated the time of maximal activity within each trial with the animals’ reaction times, focusing on the 0.5–2 s post-stimulus window to exclude visually-triggered onset responses. In both the predictable and random hit conditions, the start time of the intrinsic activity strongly correlated with subsequent reaction times - a relationship also observed in raw data when considering the maximal activity of the visually responsive cluster (Fig. 4h, Suppl. Fig. 7a). Interestingly, these changes were contingent on the presence of the stimulus, as misses close to the offset of the stimulus did not show any enhancement (Suppl. Fig. 7b). These findings suggest that feedforward sensory drive recruits a specific mode of intrinsic activity – blobs – that act as a signal enabling task engagement.

To verify if blobs specifically enhanced the visually responsive neuronal activity, we used a similarly dim, but smaller visual stimulus that is close to the perceptual limit of mice ^39^ (2° visual angle), which matches the average size of a blob. In individual trials, we observed a marked increase in activity near the time of stimulus presentation for hit relative to miss trials (Fig. 4i). This observation corresponded to an increase in the overall number of neurons that were recruited in hit compared to miss trials in the expert (but not in the beginner) stage (Fig. 4j). Consistent with this finding, NMF analysis of expert sessions revealed a marked increase in blob occurrence at and near the stimulus location (as defined in Fig. 3d,g) only during hit trials (Fig. 4k,l). Remarkably, the visual responsive cluster was not a good predictor of reaction times with the small stimulus, but the emergent activity surrounding it was (Fig. 4m). Together, these results indicate that blob-like dynamics are a key phenomenon during attentive behavior: increases in blob recruitment track changes in the attentive state. When coincident with stimulus presentation, blobs appear to enhance behavioral performance, indicating that this mode of activity captures both task outcomes and moment-to-moment fluctuations in attentional focus.

### Intrinsic dynamics can arise through local excitatory-inhibitory interactions

The pronounced modulation of intrinsic dynamics across attentive states raises the question of the underlying mechanism. Because blob-like features are uncorrelated with behavior, reward, or stimulus location, they are unlikely to arise from structured topographic input, and instead may emerge from local circuit interactions. To test whether this is plausible, we computationally screened many minimal SC network models that generalized our previously proposed, non-reciprocal feedback-driven Ising framework ^40^. Here, complex collective dynamics emerge from the interplay of several mechanisms, whose relative strength is determined by five biologically interpretable parameters (Fig. 5a): intrinsic spiking threshold, local excitatory coupling, the strength and spatial extent of the inhibitory surround, the decay timescale for circuit inputs, and the overall excitability (*β*, analogous to the “inverse temperature” of statistical physics). In this Ising-like paradigm, the dynamics of the binary (active / inactive) spatial units of the model – which correspond to activity voxels in Figs. 2, 3, and 4 – are given by stochastic state updates performed during successive Monte Carlo simulation sweeps.

**Fig. 5.**
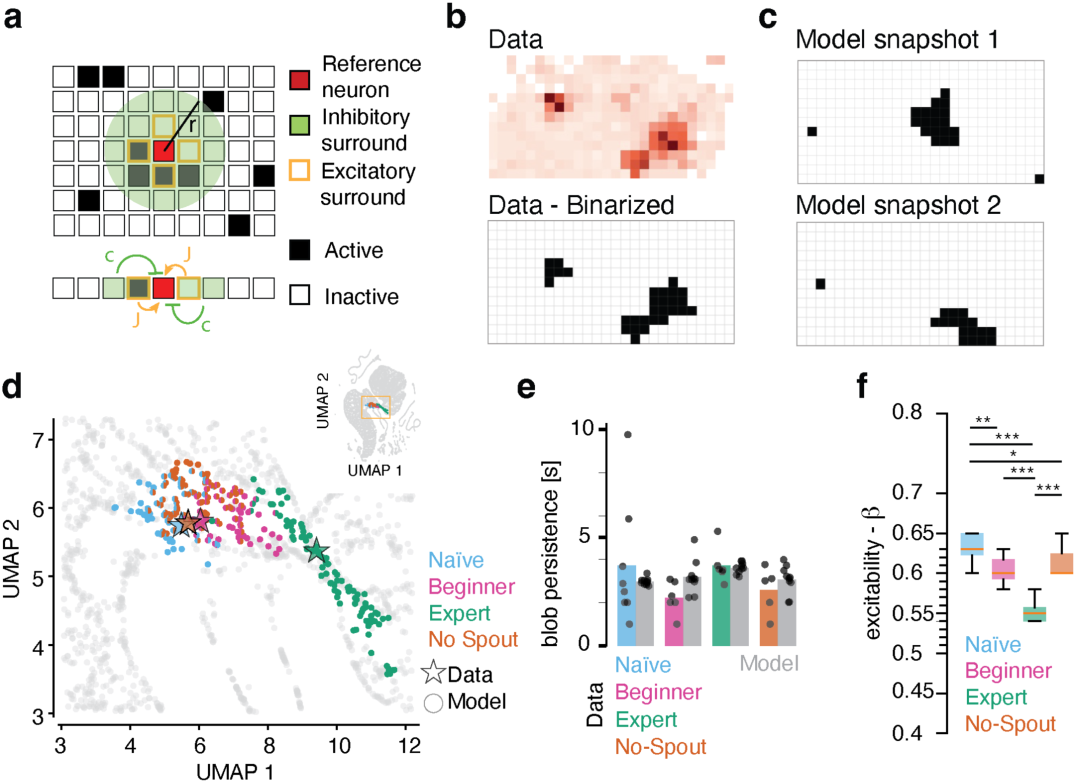
A minimal circuit model captures intrinsic dynamics and state transitions. **a.** Schematic of the model. Binary units on a 2D spatial lattice interact via nearest-neighbor excitation (*J*) and surround inhibition (radius *r,* strength *c*). **b.** Top: rasterized population activity from an example recording. Bottom: thresholded activity yields a binary spatiotemporal representation comparable to the model output. **c.** Representative model snapshots from best-matching parameter sets reproduce blob-like dynamics in the data. **d.** Zoom in on a 2D UMAP embedding of ∼25,000 models (inset) with randomly sampled parameters; colored points indicate the top 100 matches per condition (legend). **e.** Best-matching models recapitulate temporal blob persistence. **f.** Best-matching *β* is modulated by attentive state and condition.

To match the models’ output to data, we binarized the measured per-voxel neural activity by thresholding (see Methods; Fig. 5b, Suppl. Fig. 8). Thresholding preserved the essential spatiotemporal organization of the population activity across conditions and within sessions. We then performed an extensive computational screen across ∼25 000 parameter sets for our model. To align the model’s arbitrary time step to the real (experimental) time, we calibrated the decay timescales of the total activity autocorrelation functions in the model and the data (see Methods; Suppl. Fig. 9c–f). Representative snapshots of simulated activity recapitulated the experimentally observed blob-like features (Fig. 5c) for parameter sets closely matching data. Data-model match was quantified by the Wasserstein distance between the predicted and the measured distribution of instantaneous neural activity as well as the measured occurrence distribution of blob-like spatial features. The Wasserstein distance allowed us to visualize and interpret the multitude of computationally-screened models, jointly with the data, in the UMAP projection, and to identify ensembles of best-matching models for each condition (Fig. 5d; see Methods; Suppl. Fig. 9a, b, g, h).

Best-matching models recapitulated the temporal blob persistence (Fig. 5e) – an emergent dynamical timescale beyond what was directly fitted – in a spatially unbiased manner (Suppl. Fig. 9k–n). While multiple parameter sets reproduced the distributional features of the data (Suppl. Fig. 10a–d) and revealed trade-offs between the inhibitory strength and its spatial range (Suppl. Fig. 10e,f), the neural excitability *β* (Fig. 5f) robustly emerged as a single global parameter that had to change to match activity patterns across different attentive states. Varying *β* can thus be seen as a “dial” that can be precisely tuned, with small fractional changes mimicking transitions between different attentive states; larger changes drive qualitative shifts from fully asynchronous activity to the spatiotemporally correlated, large-scale, blob-like features, all without requiring any spatially structured or retinotopic input. A biologically plausible interpretation is that *β* reflects the influence of global neuromodulatory inputs, which could flexibly regulate circuit excitability and thereby control transitions between attentive states.

### Minimal circuit model recapitulates state-dependent amplification of visual responses

We next included visual stimulation into our model and compared the predicted responses with our experimental observations. To calibrate the visual input in the model, we tuned the local bias applied to the stimulated patch, for each of the 10 best naïve-matching models, so that the models reproduced the visual response amplitude observed in naïve animals. This input consisted of two brief pulses, at stimulus onset and offset, delivered to a spatially localized set of voxels matching the size and position of the experimentally stimulated region (Fig. 6a). The same calibrated parameters were then used to assess how strongly visual responses were amplified in the 10 best models matching either the expert or the no-spout condition (Fig. 6b). Mimicking the experimental results, visual responses were increased by recruitment of neighboring voxels, amplifying (even in the binarized data) the expert responses 3-fold (Fig. 6c, d). We used the model to quantify the stimulus strength required to induce blob emergence in 50% of trials across conditions. The effect was striking: whereas the expert state readily supported the recruitment of blob-like activity even by weak stimuli, other conditions required up to an order-of-magnitude stronger input to evoke comparable large-scale intrinsic events (Fig. 6e). Notably, the recruitment of intrinsic activity closely mirrored behavioral performance in the experimental data. In binarized recordings obtained with small visual stimuli, hit trials recruited blob-like events that were largely missing in miss trials and the no-spout condition (Fig. 4i, 6f,g, left). These changes matched the transition from expert to no-spout regimes observed in the model (Fig. 6g, right). Together, these results indicate that blob-like activity can emerge intrinsically from local circuit interactions while being flexibly gated by weak sensory input. This regime resembles systems poised near a critical point, where small perturbations can trigger large-scale responses ^41^, and suggests a mechanistic link between global neuromodulatory control, intrinsic dynamics, and the selective amplification of behaviorally relevant sensory inputs (Fig. 6h).

**Fig. 6.**
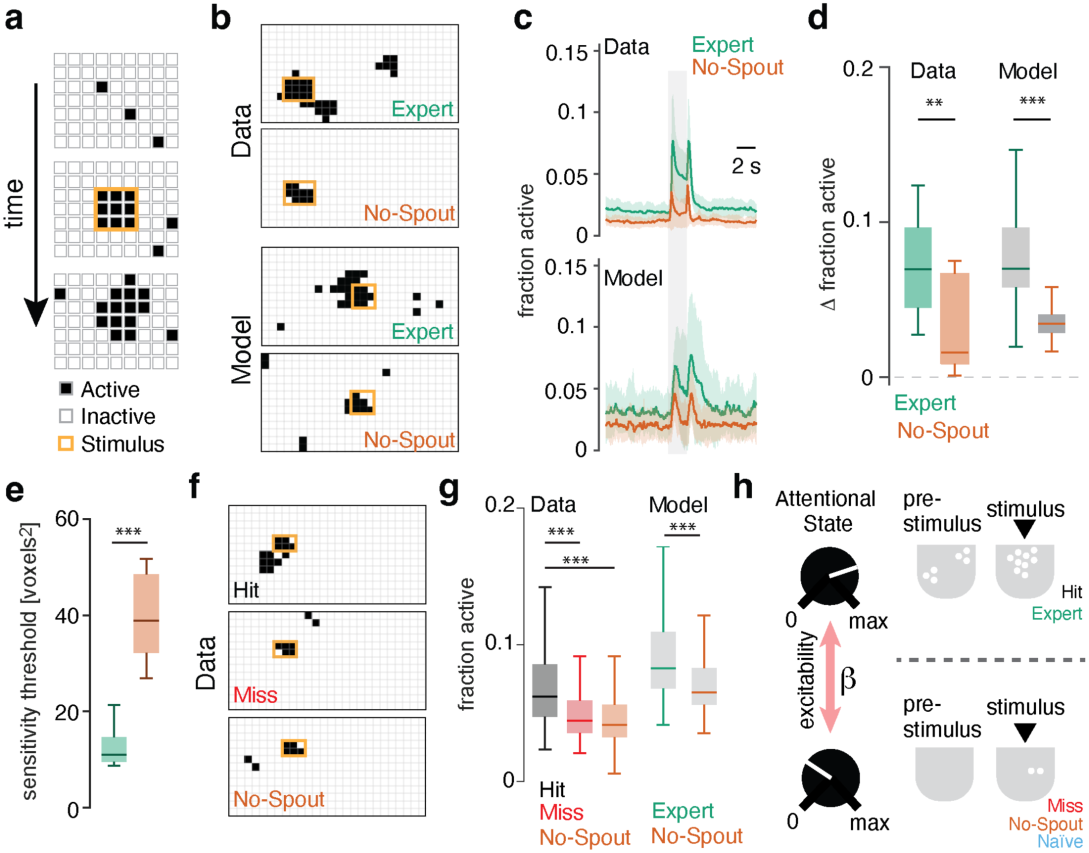
State-dependent amplification of weak sensory input is generated by local circuit interactions. **a.** Model schematic. A localized patch of “stimulus-driven” cells (orange square) was transiently stimulated by increasing the local bias before being released back to baseline dynamics. **b.** Representative stimulus-period activity frames from binarized experimental data and matched simulations under Expert and No-Spout conditions. **c.** Average temporal dynamics across the full field of view corresponding to the conditions shown in (**b**). **d.** Quantification of stimulus-evoked response changes shown in (**c**). **e.** Sensitivity threshold, defined as the stimulus size producing amplification in 50% of trials. **f.** Representative stimulus-period activity patterns for hit, miss, and no-spout conditions from binarized experimental data using a small stimulus. **g.** Quantification of activity patterns evoked by a small stimulus in (**f**) and comparison with model responses to stimuli of matched size. **h.** Summary schematic. Superior colliculus shifts intrinsic dynamics across state- and attention-dependent excitability regimes, putatively through modulation of β, a proxy for neuromodulatory gain control. All statistical details are provided in Suppl. Table 1.

## Discussion

Efficient sensory processing depends on the flexible allocation of neuronal resources to task-relevant computations. Crucially, the relevance of visual input is not fixed, but varies with context ^42^ and expectation ^43^. This adaptive prioritization is captured by saliency map models ^44^, which posit a continuous evaluation of the visual scene to highlight behaviorally relevant locations. Such maps result from the interaction of bottom-up features – such as contrast, color, and orientation ^44^, which can be approximated by deep neural networks ^45^ – with top-down influences reflecting task demands, expectations, and internal goals ^46^. In primates, including humans, saliency map activations are thought to guide subsequent gaze shifts, providing an overt readout of attention ^7^.

The superior colliculus (SC) appears to be a key hub in this integration, transforming retinal input into goal-directed behavioral commands, including those underlying visuospatial attention ^7^. Neurons in the superficial SC (sSC) integrate bottom-up retinal signals that are known to represent streams of color, contrast, or orientation ^47–49^ in a largely non-selective manner ^28,50,51^, whereas deeper layers combine visual information with other sensory modalities to generate a motor map ^30,52^. Anatomical and histochemical analyses of the sSC reveal extensive modulatory projections ^24–27,53^, suggesting the presence of non–stimulus-driven activity that could influence SC dynamics. Consistent with this view, modulation of SC neuronal activity has recently been implicated in selective visual attention ^54,55^, primarily through changes in perceptual sensitivity and response selection, rather than through shifts in the decision criterion ^56^. Despite these advances, the mechanisms that give rise to such dynamic processes remain poorly understood. Intrinsic neural fluctuations, potentially reflecting prior expectation ^3,4^, task demands ^5^, and attentional focus ^6^, often appear indistinguishable from noise. Moreover, because vision operates on a moment-to-moment basis, linking the directly observable rapid neuronal dynamics to stable attentive states and behavioral performance remains challenging.

Here we identify and characterize a new class of intrinsic population dynamics in the SC. These dynamics support attentive, goal-directed behavior, defining a network state in which behaviorally relevant stimuli recruit additional internally generated activity, thereby amplifying responses relative to unattended or disengaged conditions (Figs. 1-4). These novel dynamics are comparable in magnitude to visually evoked responses, yet sample the retinotopic map in an apparently stochastic manner (Fig. 2), rendering them invisible in trial-averaged activity and thus easily dismissed as noise (Figs. 1, 4). Strikingly, however, they reliably predict both the attentional state (Fig. 1) and the behavioral performance (hit versus miss) on a trial-by-trial basis, even before stimulus onset (Fig. 4). The emergence of this state-dependent intrinsic activity (Fig. 5), and its dynamic recruitment by sensory input (Fig. 6), can be explained by local excitatory–inhibitory interactions within the superior colliculus, giving rise to a dynamic saliency map. In the resulting framework, internally generated dynamics interact with sensory input to create “salience” – in mechanistic terms, to create neural representations that prioritize behaviorally relevant information. Notably, we observe such dynamics in the SC, a circuit specialized for linking sensory signals to behavioral priorities and actions.

Although neuronal correlates of learning have been linked to the SC ^57^, the changes we report appear to be state- rather than learning-dependent, reverting immediately to the pre-learning state upon removal of the licking spout (no-spout condition). Sensory responses are likewise enhanced on hit compared to miss trials (Fig. 4a–d), paralleling the reports in the somatosensory cortex ^58^, and recapitulating the differences between expert and no-spout states (Fig. 5). When stimulus location is predictable (block presentation), responses are rapid and robust, consistent with immediate recruitment of intrinsic activity. By contrast, when stimulus location is unpredictable (random presentation), initial visual responses are weaker and followed by delayed, spatially structured activity (Fig. 4f-g). In both conditions, the intrinsic, blob-like activity that we characterized correlates with reaction time (Fig. 4h) independently of impulsivity, as false-alarm rates are comparable across conditions (Suppl. Fig. 1l). Blobs are thus a signature of a robust, intrinsic neural population state that amplifies weak sensory signals in an attention-dependent manner directly linked to goal-directed behaviors. Taken together, these observations bear all the hallmarks of a neural substrate for functions previously attributed to the SC ^56^.

Since blobs resemble localized spotlights of activity that emerge at varying locations across the visual field, one possibility is that they reflect a continuous sampling process that transiently probes the sensory input for potentially relevant stimuli. Unlike the classical attentional spotlight models, blobs do not become spatially biased before the appearance of a behaviorally relevant stimulus (Fig. 3). Instead, they appear to define a permissive network state in which weak sensory inputs can recruit nearby intrinsic activity. In this framework, blobs do not encode attentional priority by themselves; rather, they provide a dynamic substrate with which the sensory inputs interact to generate saliency-like representations via additional blob recruitment when stimuli become behaviorally relevant (Fig. 3 & 4). These stimulus-locked blobs effectively instantiate a “dynamic saliency map”, a behaviorally-relevant representation that can guide subsequent actions. Thus, rather than spatially focused attentional phenomena, blobs reflect a global, preparatory network state that can increase the sensitivity to sensory signals. In particular, these SC dynamics amplify visual responses irrespective of the stimulus valence, suggesting that valence-specific modulation may arise from additional brain areas, such as visual cortex ^22^ or parabigeminal nucleus ^59^. Both regions are known to influence SC activity and have been implicated in visuospatial attention ^60^.

How are these state-dependent dynamics generated and regulated within the SC circuit? Our modeling work identifies a balance between intrinsic excitability, local excitation, and surround inhibition as the minimal substrate required to generate the observed intrinsic blob-like dynamics. Modulation of this balance—captured by changes in overall excitability β—acts as the key control parameter governing transitions between network states (Fig. 5, 6h). Mechanistically, our modeling suggests that *in vivo* neuromodulatory inputs could flexibly gate the emergence of intrinsic activity (Fig. 5) and the amplification of visually evoked responses, thereby enhancing weak sensory signals (Fig. 6). This interpretation is supported by the dense neuromodulatory innervation of the superficial SC ^23–27^ as well as by recent evidence that attentional engagement is accompanied by increased excitability in the intermediate SC prior to behavioral responses ^55^. Together, our model provides a mechanistic framework linking neuromodulatory control across cellular excitability, circuit dynamics, and behavior ^31^.

Recently described intrinsic activity patterns in the midbrain of fish ^61,62^ and birds ^63^ bear a striking resemblance to the activity patterns reported here. For example, in the zebrafish larva tectum, structured spontaneous activity mirrors visually evoked responses to prey, producing “hallucinations” that can trigger isolated tail movements ^61^. In the avian tectum, brief flashes of light elicit structured spontaneous events that enhance the saliency of visual stimuli and are therefore well positioned to influence spatial attention ^63^. These seemingly random spontaneous activity patterns could represent a conserved mechanism through which animals interact with the visual world on a moment-to-moment basis.

Trial-averaged responses can obscure many critical aspects of the underlying neural dynamics. Rather than being discounted as noise, trial-specific intrinsic dynamics could constitute a core feature of SC function, continuously modulating sensory processing, attentive state, and behavioral engagement. We identify a collective neural population phenomenon that fits this bill: intrinsic and sensory-driven neural activity interact to generate a dynamic saliency map. By doing so, the activity in the SC bridges the traditional divide between top-down and bottom-up accounts of attention.

## Materials and Methods

### Animals

Animal protocols were reviewed by the institutional preclinical core facility at IST Austria. All breeding and experimentation were performed under a license approved by the Austrian Federal Ministry of Science and Research in accordance with the Austrian and EU animal laws (BMF-66.018/0017-WF/V/3b/2017). During the experimental phase, mice were housed individually in standard macrolon cages with red plastic houses, running wheels and enrichment consisting of wood chips and nesting material at an inverted 12-hour light cycle. Experiments were done during the dark phase of the light cycle.

We used Rorb-IRES2-Cre mice (JAX Strain #023526) crossed with Ai148D reporter mice (JAX Strain #030328) to achieve high-level transgenic expression of the calcium indicator GCaMP6f in a Cre-dependent manner. In the superior colliculus, Rorb-Cre labels nearly all stellate cells, approximately half of narrow-field cells and horizontal cells in the superficial layers, while sparing wide-field cells^36,37^.

A total of n = 17 mice (6 F, 11 M) were used in this study, including n = 9 mice for establishing the behavioral paradigm and collecting behavioral-only data. Two-photon imaging was performed on n = 8 of these mice (mean age 14.1 ± 2.9 weeks at surgery).

## METHOD DETAILS

### Statistics

No statistical method was used to predetermine sample size. Sample sizes were chosen based on values typically used in similar studies in the field. Data collection and analysis were not performed blind to the conditions of the experiments. Imaged neuronal ROIs were included based on quality and response criteria as described below. Recordings with motion artefacts, imaging artefacts, or incomplete behavioral sessions (<30 trials) were excluded. Neurons with fluorescence signal-to-noise ratio (SNR) < 5 were excluded from analysis.

Statistical analyses were performed using custom MATLAB (MathWorks) and Python scripts. Non-parametric tests were used throughout and are specified in the figure legends and Supplementary Table 1. Data are presented as mean ± s.e.m., unless otherwise stated. Effect sizes (Hedges’ g) are reported where applicable. Significance levels are indicated as *p <0.05, ** p < 0.01, *** p < 0.001.

### Cranial window implantation surgery

Cranial window implantation was performed as described previously ^64^. In brief, mice were injected with meloxicam (20mg per kg body weight, s.c., 3.125 mg ml^−1^ solution) and dexamethasone (0.2 mg kg^−1^ body weight i.p., 0.02 mg ml^−1^ solution). Anesthesia was induced by 2.5 % isoflurane in oxygen in an anesthesia chamber and maintained at 0.7 % to 1.2 % in a stereotaxic device (Kopf), while body temperature was controlled by a heating pad to 37.5 °C. After exposing and cleaning the cranium, a 4 mm circular craniotomy was drilled above the left SC, the dura mater was removed, the left transverse sinus was sutured twice with 9-0 monofil surgical suture material (B. Braun) and cut between the sutures. Cortical areas covering the left SC were aspirated with a cell culture vacuum pump (Accuris), and a 3 mm circular coverslip, glued (Norland optical adhesives 61) to a stainless-steel conical ring, was inserted with the glass flush on the surface of the SC. After filling the surrounding cavity with Dura-Gel (Cambridge Neurotech) the insert was fixed in place with VetBond (3M). Finally, a custom-designed TiAl_6_V_4_ head-plate was affixed to the cranium by sequential application and curing of (1) All-in-One Optibond (Kerr), (2) Charisma Flow (Kulzer), and (3) Paladur (Kulzer). Mice were given 300 µl of saline and 20 mg kg^−1^ body weight meloxicam (s.c.) before removing them from the stereotaxic frame and letting them wake up while keeping them warm on a heating pad. Another dose of 20 mg per kg body weight meloxicam s.c. and 0.2 mg per kg body weight i.p. dexamethasone was further injected 24h after the conclusion of the surgery. After the implantation surgery, mice were allowed to recover for at least one week.

### Behavioral Paradigm

Visual spatial attention task. Mice were trained on a classical conditioning visual detection task. Head-fixed animals were positioned on a custom-designed spherical treadmill (20 cm diameter) in front of a panoramic dome projection system (80 cm diameter). Mice were trained to detect a white square (∼8° visual angle) that appeared at one of several retinotopic locations, including the central binocular zone (0° azimuth) and lateral monocular zones (± >65° azimuth, adjusted based on imaged retinotopic field). All stimuli were positioned in the dorsal visual field, approximately 20° above the equator. In a subset of experiments, a smaller stimulus (2° visual angle) was used. Each trial consisted of a variable wait period (8-13 s) during which mice were required to suppress spontaneous licking, followed by a 2 s stimulus and reward window. The first lick during this window triggered reward delivery (∼ 350 ms after the lick). Extraneous licks during the wait period were discouraged with a timeout and mild auditory penalty. Specifically, timeouts were triggered when the lick rate exceeded a threshold of 0.2 licks per second.

Motivation, reward delivery and lick detection. Instead of water restriction, mice had free home-cage access to 2 % citric acid water ^65^, which reduced voluntary water intake while preserving motivation for a palatable liquid reward during training. The reward consisted of Humana SL soy-based, milk-free baby formula and was delivered through copper reward spouts, insulated on their outer surface to avoid electrical cross-talk with the lick sensor. Licks were detected by capacitive sensing on the spout using an Adafruit MPR121 12-key capacitive touch-sensor breakout (Adafruit #1982), and reward was dispensed by a custom-built syringe pump. Lick detection and pump actuation were driven by the same Arduino microcontroller, ensuring tight temporal coupling between the first detected lick and reward onset.

Learning stages. Animals were classified into distinct stages: naive (before training), beginner (first training session), expert (high hit rate coupled with suppression of non-task-related licking, typically achieved after ∼10 sessions), and no-spout (same as expert animals, but prevented from engaging in the task by removing the reward spout). Animals remained engaged for up to 150 trials per session, with reaction times sharply increasing beyond 120 trials. Sessions were therefore terminated at that point to ensure similar levels of engagement across trials. Training progression was managed by a custom MATLAB script that advanced animals through a graded sequence of difficulty levels (from initial lick shaping to full detection with penalties), with per-session parameters (timeout duration, lick threshold, stimulus contrast) adjusted adaptively based on each animal’s running hit rate and false-alarm rate. Session-to-session state (current level, adaptive parameters, performance history) was saved per animal, enabling longitudinal training with consistent criteria across sessions.

Stimulus conditions. Visual stimuli consisted of a dim square that covered ∼8 degrees of visual angle. Three stimulus positions (P1, P2, P3) were arranged symmetrically about the animal’s visual midline on a screen placed 38 cm from the eye. P1 was chosen individually per animal at the retinotopic location that elicited a clear visual response centered in the imaging window. P3 was its mirror image across the midline in the ipsilateral hemifield, and P2 lay exactly between P1 and P3 (at the midline, directly in front of the animal). P1 and P3 were therefore at equal and opposite azimuths, with P2 at 0°. In a subset of experiments, a smaller stimulus (2 degrees of visual angle) was used, which approximately matched the average size of intrinsic activity blobs and prevented occluding their presence with visually driven responses. Expert animals were tested in separate sessions with two stimulus presentation regimes. In predictable sessions, the stimulus appeared at a fixed retinotopic location, creating an internal spatial expectation. The location was changed every 25-30 trials to a new fixed position with these subsessions allowing comparison of attention-dependent modulation across retinotopic space. In random sessions, the stimulus location varied from trial to trial, increasing task difficulty level and requiring the animal to distribute attention across multiple spatial positions.

### Head-fixed *in vivo* recordings

Imaging experiments were performed on a custom-built system. In short, mice were head-fixed while awake using a custom-manufactured clamp (for imaging: connected to a three-axis motorized stage (8MT167-25LS, Standa) and could run freely on a custom-designed spherical treadmill (20-cm diameter). Running behavior was recorded by a pair of ADNS-3080 (iHaospace, Amazon) optical flow sensor modules, focused with 25 mm lenses (AC127-025-AB-ML, Thorlabs) on a small patch at orthogonal locations of the Styrofoam ball and illuminated by an 850 nm LED. The alternating sensor readout was controlled at 50 frames/s by an Arduino Uno running custom scripts. The 4 signal channels from the sensor were linearly mapped to movement speed in the forward, sideways and rotational axes based on regular calibration with synchronous measurement of image translations and rotation at the ball’s apex. Eye and body movements were recorded at 50 fps with infrared illumination (850 nm) with a Camera (acA1920-150um, Basler) and an 18-108 mm macro zoom objective (MVL7000, Thorlabs), pointed at the right side of the mouse via an infrared mirror. Eye position and saccades were determined post-hoc as previously reported ^64^, by first labeling eight points around the pupil with DeepLabCut ^66^, which were fitted to an ellipse, and the center position was transformed to rotational coordinates. Fast eye position changes of more than 45 ° s^−1^ and at least 3 ° amplitude on a 0.7 s median filtered trace were defined as saccades. The ellipse area in mm^2^ was determined as pupil size.

Visual stimuli were projected by a modified LightCrafter (Texas Instruments) at 60Hz (DLP Lightcrafter evaluation module, Texas Instruments), reflected by a quarter-sphere mirror (Modulor) below the mouse and presented on a custom-made spherical dome (80cm in diameter) with the mouse’s head at its center. For imaging experiments, a double bandpass filter (387/480 HD Dualband Filter, Semrock) was positioned in front of the projector to minimize light contamination during imaging. In both setups, the blue LED in the projector was replaced by UV (LZ1-00UB00-01U6, Osram) and in addition, in the multi-photon setup, the green LED was replaced by a cyan LED (LZ1-00DB00-0100, Osram) not to interfere with the Calcium imaging wavelengths. The reflected red channel of the projector was used for synchronization and captured by a trans-impedance photo-amplifier (PDA36A2, Thorlabs) and digitized. Stimuli were designed and presented with Psychtoolbox-3 ^67^, running on MATLAB (MathWorks) on Microsoft Windows 10 systems. Stimulus frames were morphed on the GPU using a customized projection map and an OpenGL shader to counteract the distortions resulting from the spherical mirror and dome. In both setups, the dome allows the presentation of mesopic stimuli from ∼100 ° on the left to ∼135 ° on the right in azimuth and from ∼50 ° below to ∼50 ° above the equator in elevation.

### *In vivo* Calcium imaging

Two-photon imaging in superior colliculus was performed as described previously ^64^. In brief, ScanImage (Vidrio Technologies) on MATLAB 2020b (MathWorks) controlled a custom-built microscope using a pulsed Ti:Sapphire laser (Mai-Tai DeepSee, Spectra-Physics) set at wavelengths between 920 and 950 nm. The beam was expanded to underfill the back-aperture of the objective (×16 0.8-NA water-immersion, Nikon) and scanned through the tissue by a galvanometric-resonant (8kHz) mirror combination (Cambridge Scientific) and a piezo actuator (P-725.4CA, Physik Instrumente) controlling the objective. Emission light was measured with GaAsP photomultiplier tubes (H10770B-40, Hamamatsu) following collection by a dichroic mirror (FF775-Di01, Semrock) and channel splitting (580nm long-pass, FF580-FDi01, Semrock) as well as filtering (green: FF03-525/50; red: FF01-641/75; Semrock). The signals were then amplified by a TIA60 amplifier (Thorlabs) and digitized with a PXI system (PXIe-7961R NI FlexRIO FPGA, NI 5734 16-bit, National Instruments). Average laser output power at the objective ranged from 100 to 125mW. A field of view (FOV) of 0.72 – 1.51 mm^2^ (median of 0.82 mm^2^) was imaged over 2–5 planes (median 2 planes) with a plane distance of 40 µm at a pixel size of 1.18 – 1.57 µm (median of 1.25 µm) and a volume rate of 5–10 Hz (median of 10 Hz).

### *In vivo* Calcium imaging analysis

Imaging data was motion corrected and ROI segmented with suite2p (v0.10.0)^68^ followed by a manual curation step based on morphological and activity shape. Further analysis was performed in MATLAB (MathWorks). dF/F_0_ was estimated as previously ^43,64^, by subtracting neuropil contamination with a factor of 0.5, defining F_0_ baseline as the 8th percentile of a moving window of 15 s ^69^ and finally subtracting and then dividing the fluorescence trace by the median of the same 15 s window. The fluorescence signal-to-noise ratio was defined for each ROI by dividing the 99^th^ percentile of the dF/F_0_ trace (‘signal’) by the standard deviation of its negative values after baseline correction (‘noise’). Only neurons with a fluorescence SNR ≥5 were included in further analysis.

#### Visual Response Classification

Visually responsive neurons were classified using k-means clustering with *k = 2* (responsive vs. non-responsive), 100 replicates with different initializations, and Euclidean distance metric. Clustering was performed on concatenated z-scored activity traces across all trials, using a time window from 0.1 to 2.4 seconds after stimulus onset. For each recording, the smaller cluster was assigned as the visually responsive population, as it represented the specific subset of neurons recruited by the localized stimulus. This approach enabled unbiased identification of stimulus-driven cells across learning stages and conditions.

#### Population Entropy

Population entropy (PE), a measure of how evenly neural activity is distributed across the imaged population at each timepoint, was computed as the Shannon entropy of the normalized activity distribution:

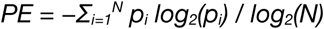

where *p_i_ = (X_i_ + ε) / Σ_j_(X_j_ + ε)* represents the normalized activity of neuron i, X_i_ is the raw activity (clipped at zero), a small constant of *ε = 10^−8^* prevents numerical errors from *log(0)*, and *N* is the total number of neurons. Normalizing by *log_2_(N)* yields PE values between 0 (all activity concentrated in one neuron) and 1 (activity evenly distributed). Low PE instances indicate spatially-clustered activity patterns.

#### Entropy drop detection

Entropy drops were identified using peak detection on the inverted entropy trace with MinPeakHeight = - 0.8 and MinPeakProminence = 0.05. Entropy-triggered averaging was performed by aligning activity to detected low-PE timepoints.

#### Spatial extent of blob events (FWHM)

To quantify the spatial extent of intrinsic and visually evoked activity patterns, we computed activity profiles centered on a reference voxel and measured the full-width at half-maximum (FWHM). Two approaches were used. For intrinsic dynamics (Fig. 2l,m), single-frame maps at entropy drop were centered on the maximally active voxel within the respective pre-stimulus or stimulus periods. For visually evoked responses (Fig. 2n,o), trial-wise spatial maps during the stimulus onset were centered on the centroid of the visually responsive voxel cluster. Maps were averaged across trials and recordings to obtain a mean two-dimensional activity map for each condition. A one-dimensional radial profile was then computed by averaging values over concentric rings centered on the reference voxel (radial step: 40 μm, corresponding to voxel spacing). This profile was fitted with a single exponential decay using non-linear least squares, and the resulting fit was used to estimate the FWHM.

#### Coefficient of Variation Analysis

Response variability was quantified using the coefficient of variation (CoV):

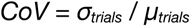

where *σ_trials_ and μ_trials_* are the standard deviation and mean of single-trial responses across trials for each neuron. CoV was computed for different time windows: pre-stimulus baseline and stimulus epoch.

#### Grid-Based Spatial Activity

For spatial analysis, the imaged region was discretized into a 13 x 26 voxel grid (each voxel measuring 40 x 40 x 250 µm^3^). Only voxels containing more than five neurons were included. The activity of neurons within each voxel was averaged for subsequent analyses.

#### NMF Decomposition

To characterize the spatial structure of intrinsic dynamics, we decomposed the data with non-negative matrix factorization (NMF) using cross-orthogonality and L1-sparsity regularization to extract distinct, non-overlapping components. We used the seqNMF implementation^70^ with the temporal factor length set to L = 1. This disables the temporal convolution, so each component is a single time slice and the decomposition reduces to ordinary NMF, V ≈ WH, where W holds the spatial components and H their temporal activation profiles. We did not use the method to discover temporal sequences, but utilized its cross-orthogonality penalties to enforce orthogonality between components in both space (W) and time (H).

Off-diagonal cross-orthogonality penalties were applied to the spatial overlap W^T^W and the temporal co-activation HH^T^ (with L = 1 the smoothing kernel S in HSH^T^ has length 2L−1 = 1, so the temporal term reduces to instantaneous orthogonality of H), penalizing similarity between different components while preserving their magnitude:

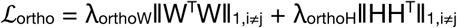

We used k = 10 components, L = 1, and 100 iterations, with regularization weights λ = 0.025 (factor-competition penalty), λ_orthoW_ = 2.5 and λ_orthoH_ = 0.5 (spatial and temporal cross-orthogonality), and λ_1W_ = 1.0 and λ_1H_ = 0.75 (L1 sparsity on W and H).

The decomposition explained ∼50% of the total variance, with similar reconstruction errors across learning stages. To isolate intrinsic dynamics, the mean visually evoked response was subtracted from each trial before applying seqNMF.

#### Overlap Analysis

Overlap between activity patterns was quantified using the Jaccard index:

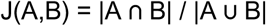

For spontaneous activity, A and B represent sets of voxels participating in two blob-like events: low indices indicate spatially variable blob positions, while high indices indicate stable, recurring spatial patterns. For stimulus-evoked responses, A and B represent sets of neurons participating in two response events: low indices indicate dynamic response compositions, while high indices indicate consistent neuron partnerships (“response fidelity”).

#### Spatial Activity Maps

Blob occurrence frequency maps were created by detecting blobs across the frames of a trial on the 13 × 26 spatial voxel, calculating each blob’s centroid using MATLAB’s regionprops, and aggregating centroid positions across trials to create spatial density maps (units: blobs/second/location).

#### Support Vector Machine (SVM) classification

Learning stage (Fig. 2k) and trial outcome (Fig. 4e) were decoded using SVM trained on single-trial population entropy (PE) in the 2 s preceding stimulus onset. For each trial, the PE trace within this window served as the feature vector and was z-scored per recording. Binary classifiers (Naïve vs. Expert, Beginner vs. Expert, No-Spout vs. Expert, and Hit vs. Miss) were implemented using MATLAB’s *fitcsvm*, whereas the four-class learning-stage classifier (Fig. 2k) used *fitcecoc* with one-vs.-one binary SVM templates. All classifiers used a Gaussian (RBF) kernel with a box constraint of 1 and standardized features (zero mean, unit variance). Performance was assessed using 5-fold cross-validated accuracy (*kfoldLoss*). Classes were balanced by random subsampling of the larger class within each fold. For the Hit vs. Miss classifier (Fig. 4e), trials were stratified by lick latency, and classification accuracy is reported as a function of reaction-time threshold, comparing miss trials of increasing latency against hits.

#### Computational Modeling

To investigate whether blob-like assemblies can arise from intrinsic SC circuitry, we implemented a 2D Ising model with local excitatory–inhibitory interactions on a 39 × 78 lattice (3× experimental voxel dimensions) with periodic boundary conditions.

#### Spin dynamics

Each lattice site i holds a binary spin σ_i_ ∈ {−1, +1} updated via the heat-bath algorithm:

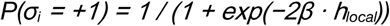

where β is the inverse temperature and the local field is:

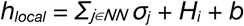

with NN denoting nearest neighbors (4-connected), H_i_ an auxiliary field, and b a bias term.

#### Auxiliary field dynamics

The auxiliary field H_i_ provides surround inhibition:

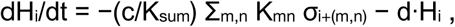

where c is the coupling strength, d the decay constant, the sum runs over the 2-D kernel offsets (m, n), and K_sum_ = Σ_m,n_ K_mn_ is the sum of the kernel weights (normalisation). K is a diamond-shaped surround-inhibition kernel of radius r:

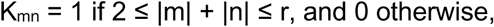

with the center (m,n) = (0,0) and the four nearest neighbors (|m| + |n| = 1) removed. These are precisely the sites already entering the local field as excitation through the Σ_j∈NN_ σ_j_ term, so that nearest neighbors contribute only to local excitation and more distant sites only to surround inhibition. The field was integrated by forward Euler with dt = 1/(L·M), so that one Monte Carlo sweep (L·M single-site updates) advances one unit of model time.

*Parameter space.* We explored 24,624 parameter combinations:

- â (inverse temperature): 19 values spanning [0.4, 0.8] with finer resolution near the critical point (0.50–0.63)
- c (coupling strength): [1, 2, 3, 4, 5, 6, 7, 8, 9]
- d (decay constant): [2, 3, 4, 5, 6, 7, 8, 9, 10, 11]
- r (inhibition range): [2, 4, 9, 13]
- b (bias): [−1.0, −0.8, −0.6, −0.4]

#### Simulation protocol

The auxiliary inhibitory field H relaxes with a characteristic timescale τ_H = (L·M)/d, where L·M is the number of lattice sites and d the decay constant (expressed in Monte Carlo sweeps). Each simulation began with an adaptive burn-in phase, the larger of 2,000 sweeps and 7τ_H (≈99.9% equilibration), followed by 100,000 recording sweeps.

#### Binarization of experimental data

To compare the experimental grid activity with continuous dF/F values to the binary spins of the Ising model, rasterized time series were thresholded at dF/F > 2, which corresponds to the top ≈ 3.9% of the per grid cell activity distribution and consistently maximizes Dice overlap with the seqNMF non-stimulus reconstruction across binarization thresholds (Suppl. Fig. 8). The resulting binary spatiotemporal patterns served as the reference for both Moran’s I and the per-frame mean-activity distribution used in matching (below).

*Spatial autocorrelation: Moran’s I.* Spatial clustering of activity was quantified using Moran’s I, computed per frame on the 13 x 26 grid (binarized experimental data) or on the 39 x 78 simulation lattice (binary spins):

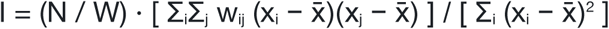

where x_i_ is the value of a given grid cell, x̄ is the spatial mean of the grid, and N is the number of grid cells. Each cell was connected to its four immediately adjacent cells (above, below, left, and right): wᵢⱼ = 1 for these neighbors and 0 for all other cells, including the cell itself. W = Σᵢ Σⱼ wᵢⱼ is the sum of all weights (the normalizing constant).

Moran’s I values range between - 1 and 1: values close to 1 represent states of perfect spatial clustering (the regime in which blob-like features dominate), values near 0 indicate no spatial autocorrelation and values near -1 are found at perfect dispersion.

To compare model and data on equal terms, simulation Moran’s I was evaluated on non-overlapping 13 × 26 tiles of the lattice. The per-position distributions were indistinguishable (Suppl. Fig. 9k–n), confirming that lattice statistics are spatially homogeneous.

#### Model-data matching

Simulated dynamics were compared to experimental data using three metrics: Moran’s I (spatial autocorrelation; above), the per-frame activity distribution (fraction of active cells), and blob persistence distribution. Combined matching used equal weights for the first two metrics, while the blob-persistence distribution was held out as a generalization check (Fig. 5e, Suppl. Fig. 9). For each Ising parameter set, the simulated and experimental distributions of each metric were compared using the 1-Wasserstein distance, approximated by the L1 distance between empirical quantile functions evaluated at 1000 equally-spaced quantiles. For each condition, the Wasserstein distances for Moran’s I and activity were z-scored across simulations and combined (0.5 / 0.5) into a single per-simulation matching score. Then the simulations were ranked, and the top 10 were selected as best-matching parameter sets (top 100 also shown in Fig. 5d / Suppl. Fig. 9a). Because the blob-persistence distribution was not used in ranking, its agreement with data is an emergent property of fits to the spatial (Moran’s I) and population-level (activity) distributions. The temporal scale factor (Monte Carlo sweeps per imaging frame) was independently calibrated per condition by matching the autocorrelation decay timescale of the total activity (Suppl. Fig. 9c–f, iߝj). This scale factor was used both to express blob-persistence values in seconds (Fig. 5e) and to set the duration of the simulated stimulus window in the perturbation experiments described below.

#### UMAP embedding for model & data visualization

To visualize how the ∼25,000 simulated parameter sets relate to the four experimental conditions (Fig. 5d, Suppl. Fig. 9a), we computed a 2D UMAP embedding (umap-learn) jointly on all simulations and conditions. The input was a precomputed pairwise distance matrix in which every entry was the 1-Wasserstein distance between the two items’ Moran’s I and activity distributions (the same metrics used for matching, above). UMAP was run with metric=’precomputed’, n_neighbors=15, min_dist=0.3, and random_state=42. All points were embedded in a single fit_transform call so that simulations and experimental conditions share the same coordinate system. The embedding is purely descriptive: parameter selection was done directly on the Wasserstein ranking, not on UMAP coordinates.

### Simulated Visual Stimulation

To test whether the model reproduces the state-dependent amplification of sensory input observed experimentally, we applied localized perturbations to the top 10 best-matching parameter sets per condition. Each perturbation trial consisted of three phases: (i) a pre-stimulus equilibration period of 400 frames during which the network evolved freely from equilibrium, (ii) a stimulus period whose duration in Monte Carlo sweeps was chosen to match the 2 s experimental stimulus window (converted via the temporal scale factor calibrated above), and (iii) a post-stimulus recovery period of 300 frames. 150 independent replicates were run per stimulus configuration, each initialized from decorrelated equilibrium states separated by at least 2τ_H Monte Carlo sweeps. The input magnitude was calibrated on the 10 best naïve-matching models so that the simulated evoked-response amplitude matched that measured in naïve animals, yielding a stimulus bias of b = +1.75. These same stimulation parameters were then held fixed and applied identically to the beginner, expert, and no-spout models, so that any difference in amplification reflects the network state (β) rather than the input. Stimulus size was varied separately (below) for the sensitivity-threshold analysis.

During the stimulus period, a centrally placed square region of the lattice received an additional local bias: the bias term b within the patch was transiently raised from its baseline value to +1.75, increasing the probability that those spins were active without forcing them. This resembles a physiological excitatory drive rather than a hard clamp. The bias was applied as a double pulse, only during the first and last frames of the stimulus period, with the network evolving freely in between, mimicking the transient on/off response of superficial-SC visual neurons. To match the experimental FOV, all metrics were evaluated on a center crop of the simulation voxel (rows 13–26, columns 26–52; 13 × 26 elements) corresponding to the experimental voxel dimensions. Stimulus sizes of 1, 2, 3, 4, 6, 8, 10, and 12 voxel elements (side length) were tested.

We tested different stimulus sizes to define how the sensitivity threshold, the size at which a stimulus is amplified 50% of the times, depends on stimulus strength (Fig. 6e).

### Quantification of stimulus-driven activity

Stimulus-evoked recruitment of additional activity was quantified as the size of the largest connected active region overlapping the stimulus site. Each frame was thresholded (dF/F > 2 for the experimental voxel; already binary for the Ising simulations) and segmented into connected components using 4-connectivity. Within the stimulus period, the largest component intersecting the stimulus region was traced across frames and its peak area (in voxel elements) was taken as the single trial response magnitude. This provided a direct readout of whether the perturbation recruited additional, blob-like responses. The identical pipeline was applied to both the experimental voxel and the top 10 best-matching simulations per condition for the Ising model (on trials matched for stimulus size). This allowed data and model comparison using the same metric, both across conditions (Fig. 6c, d) and between hit, miss and no-spout trials (Fig. 6f, g).

## Supporting information

Example (Mouse behavior) Hit/Miss

Example videos for the mean response and three single trials per condition.

Example videos for the mean response and three single trials per condition (Grid structure, Data same as Suppl. Video 1).

NMF reconstruction example videos, same as Video 3, but with the mean response subtracted (Reconstruction of data presented in Suppl. Video 2).

NMF reconstruction example videos for the mean response and three single trials per condition (Reconstruction of data presented in Suppl. Video 2).

Statistics Summary

## Data and code availability

Behavioral and functional imaging data will be deposited and made publicly available upon publication. DOIs will be listed in the key resources table. Microscopy data reported in this paper will be shared by the lead contact upon request.

All original code will be deposited and made publicly available upon publication. DOIs will be listed in the key resources table.

Any additional information required to reanalyze the data reported in this paper is available from the lead contact upon request.

## Acknowledgements

We thank Richard Krauzlis for his suggestions, advice, and for serving on FS’s thesis committee; Arjun Krishnaswamy for helpful comments on the manuscript; and Olga Symonova for technical assistance. We are also grateful to all members of the Neuroethology group for their advice, feedback, and support. This work was supported by an internal IPC grant (ISTA) to GT and MJ, the European Research Council Consolidator Grant 101086580 (MJ), and fellowships from EMBO (ALTF 1098-2017, AS) and the Human Frontier Science Program (LT000256/2018-L, AS).

## Author contributions

FS, WM, GT, and MJ designed the study. FS performed most of the *in vivo* imaging experiments. AS developed the imaging system and provided the initial description of the process. FS and AS developed the combined behavioral and imaging paradigm. WM spearheaded the data analysis pipelines and initial descriptions of the phenomenon, FS performed all subsequent data analysis. FS and AS performed the software engineering for hardware control. FS, AG and GT developed the computational model. MJ wrote the manuscript with the help of the other authors.

## Supplementary Figures

**Suppl. Fig. 1.**
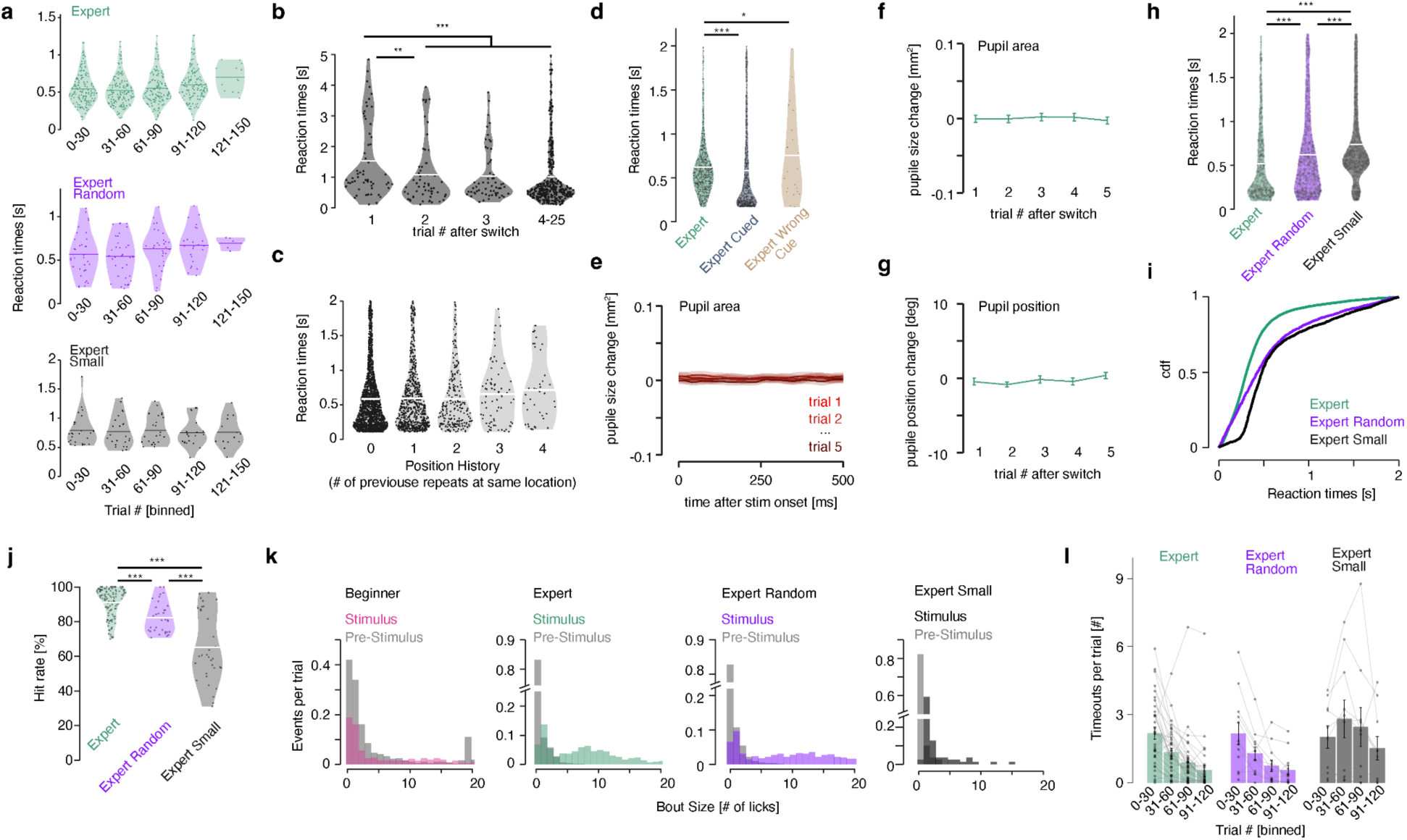
Attentional focus shifts, driven by cueing and repeated stimulation, modulate detection task performance. **a.** Reaction times (RTs) across trials for stimuli presented at a fixed location (expert), at random locations (expert random), or at a fixed location with a small stimulus (2° visual angle; expert small). **b.** RTs for stimuli presented at a fixed location for 20 trials and then switched to a novel location, aligned to the first location switch, reveal a marked RT increase immediately after the change. **c.** RTs for random-location sessions, aligned by randomly occurring repetitions at the same location. **d.** RTs for uncued, cued, and incorrectly cued stimuli show reduced or increased RTs relative to uncued trials, respectively. Animals did not respond to the cue, which consisted of a dim frame surrounding the stimulus. **e-g.** Eye tracking following stimulus onset for the predictable stimulus (**b**) show no differences in pupil dynamics (**e**), arousal level (**f**), or eye position (**g**). **h-i.** RTs for stimuli presented at the retinotopic location used for imaging, under predictable, random, and small-stimulus conditions, plotted as distributions (**h**) or cumulative density functions (**i**), reflect task difficulty. **j.** Hit rates for the conditions shown in (**h**) reveal an increase in miss rate that scales with task difficulty. **k.** Lick distributions for beginner, expert, expert random & expert small sessions, measured before and during the reward window, reveal two behavioral strategies: bursts and isolated single licks. Note, most trials have zero licks in the pre-stimulus time for experts. **l.** Timeouts triggered when lick rate exceeds a licking threshold. The threshold of 0.2 licks/s was defined for each wait period dependent on the wait-period length. Licks are accumulated during that wait period, and if the threshold is exceeded, a timeout is triggered. Timeouts decrease progressively across the session. Notably, timeouts were rarely triggered by bursts but instead by the accumulation of single licks. All statistical details are provided in Suppl. Table 1.

**Suppl. Fig. 2.**
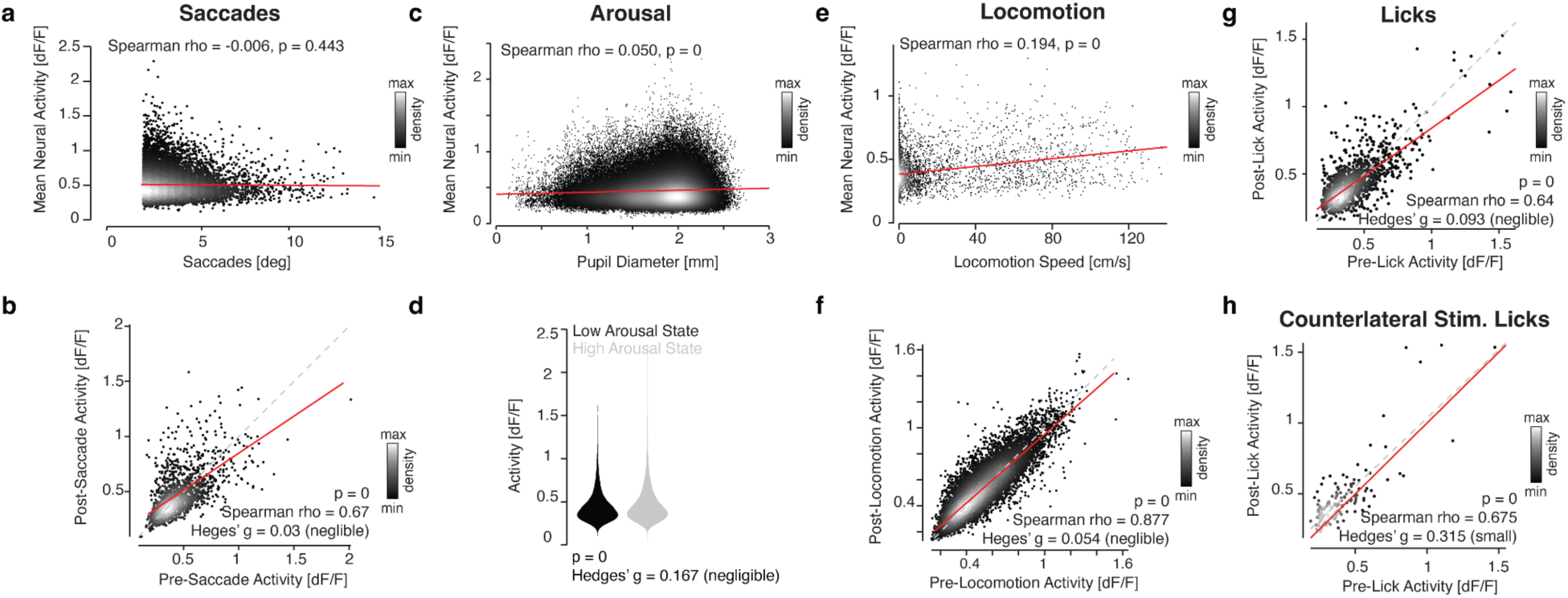
Behavioral, arousal and reward modulation don’t explain response changes in visual spatial attention. **a.** Activity changes of the entirely imaged superior colliculus are not correlated to the magnitude of saccades, (**b**) saccadic movements nor (**c**) arousal. **d.** The distribution of activities in low and high arousal states (low and high, 10% of the pupil size distribution, respectively) have negligible differences, as (**e**) correlations with locomotion speed, (**f**) the initiation of locomotion, (**g**) non-stimulus evoked licking behavior or (**h**) stimulus evoked reactions (stimulus presented contralaterally to avoid stimulus activity). Red lines: least-squares linear regressions, dashed lines: unity. All statistical details are provided in Suppl. Table 1.

**Suppl. Fig. 3.**
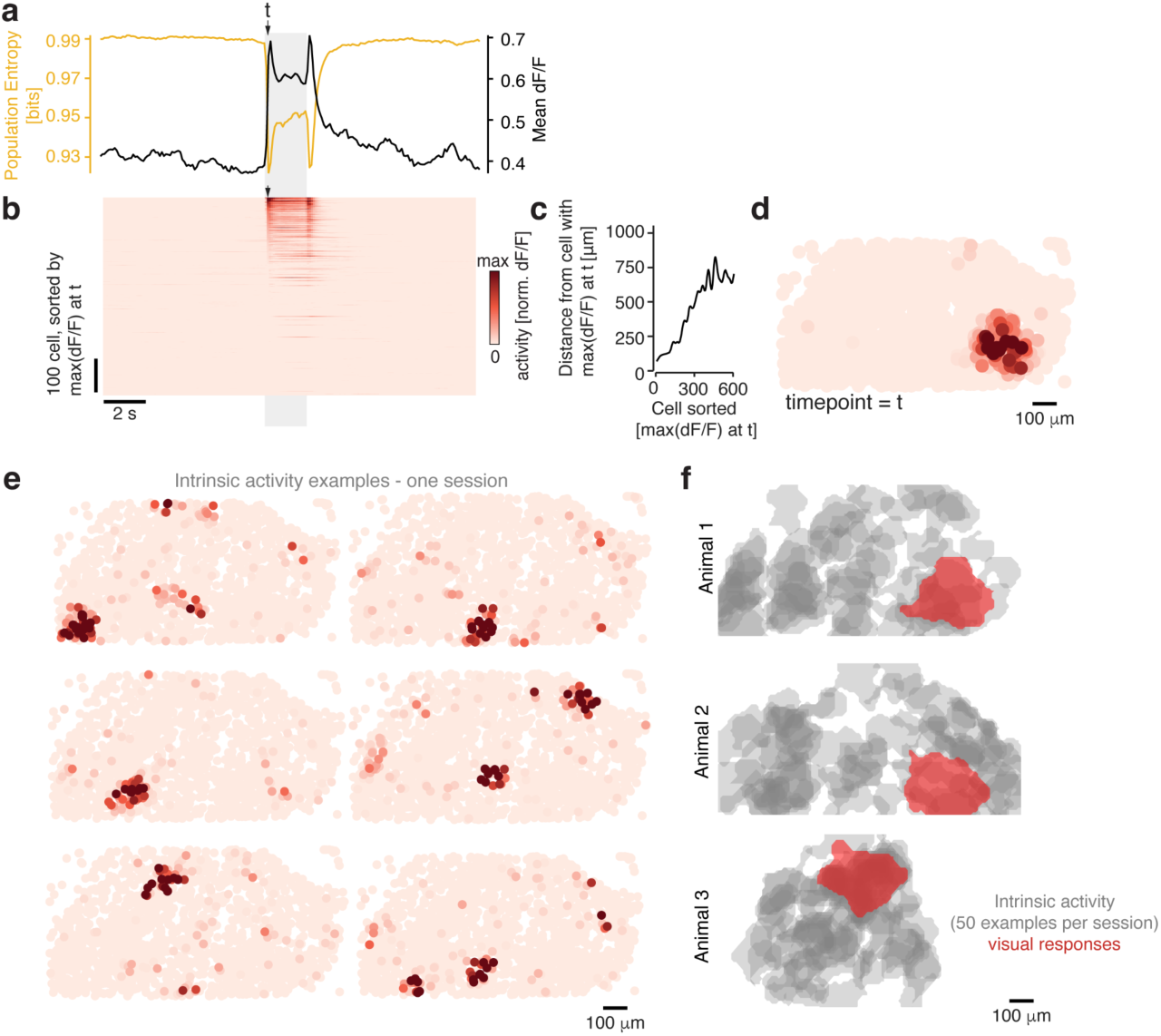
Deciphering the intrinsic activity. **a.** Representative mean session showing population entropy (PE, yellow) and mean calcium activity (black) across all trials in an expert animal. **b.** Average calcium responses from the session in (**a**), sorted by response strength at the onset of the visual stimulus (timepoint *t*, indicated by the arrowhead). **c.** Smoothed spatial distance from the maximally responsive cell, sorted as in (**b**), highlighting the spatial organization of response strength. **d.** Spatial map of mean calcium responses for each cell at timepoint *t*, revealing the visually responsive cell cluster. **e.** Example intrinsic activity events from the session shown in (**d**). **f.** Contours of 50 intrinsic activity events (grey) overlaid with the corresponding visually evoked responses (red), demonstrating a complete mapping of visual space. Data from Animal 1, same session as in (**d, e**). All statistical details are provided in Suppl. Table 1.

**Suppl. Fig. 4.**
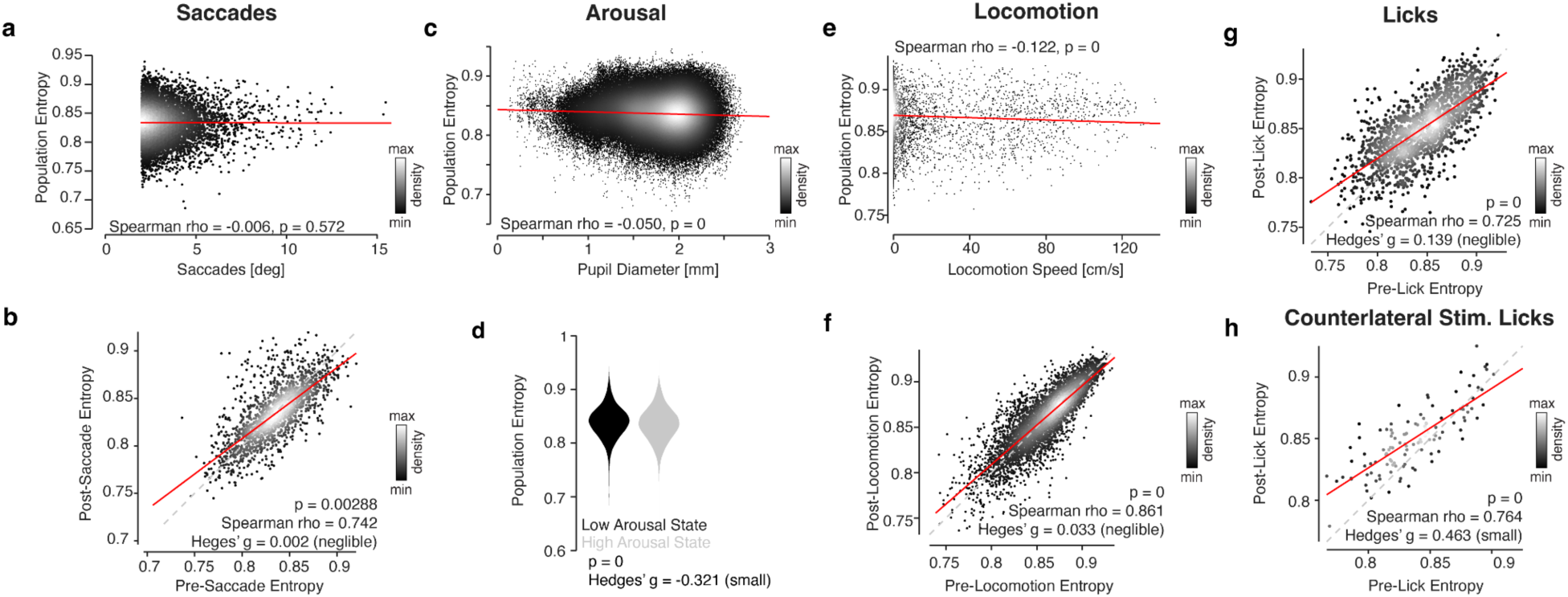
Behavioral, arousal and reward modulation do not explain response changes to population entropy. **a.** Entropy changes are not correlated to the magnitude of saccades, (**b**) saccadic movements nor (**c**) arousal. **d.** The distribution of activities in low and high arousal states (low and high, 10% of the pupil size distribution, respectively) have small differences, as (**e**) correlations with locomotion speed, (**f**) the initiation of locomotion, or (**g**) non-stimulus evoked licking behavior. **h.** Stimulus evoked licking reactions (stimulus presented contralaterally to avoid stimulus activity) show a small correlation. Red lines: least-squares linear regressions, dashed lines: unity. All statistical details are provided in Suppl. Table 1.

**Suppl. Fig. 5.**
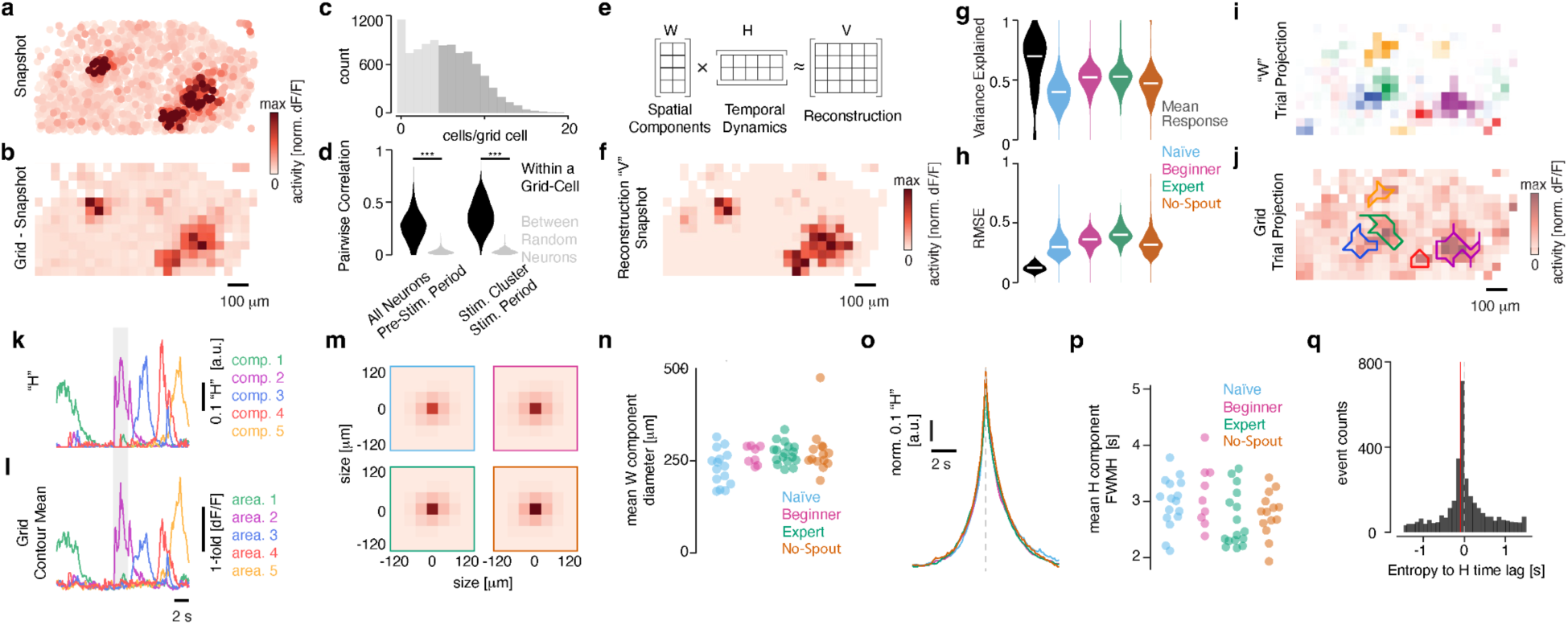
NMF decomposition with regularization penalty confirms key properties of intrinsic and collective neuronal dynamics. **a.** Representative snapshot of the spatial map of calcium responses during visual stimulation, showing a visually responsive cell cluster (center right) and two intrinsic activity blobs. **b.** Same as in (**a**), but with responses averaged across all cells within 40 μm × 40 μm voxels. Only voxels containing more than five cells were included in subsequent analyses. **c.** Distribution of cell counts per voxel. **d.** Correlations between neurons within and across voxels for all cells and visually responsive subsets, demonstrating tightly linked activity patterns that justify the voxel-based simplification of the spatial map. **e.** Schematic of non-negative matrix factorization (NMF) illustrating the spatial (W) and temporal (H) components used to reconstruct the original activity matrix (V). This factorization included a cross-orthogonality regularization penalty to minimize redundant factors and facilitate the extraction of non-overlapping neural patterns (see Methods). **f.** Example reconstruction of (**b**). **g-h.** Variance explained and the Root Mean Square Error (RMSE), computed from mean responses across conditions and experiments as well as from single trials per condition, demonstrate comparable reconstruction efficiency across all conditions. **i.** Projection of all spatial maps (W) from a single trial, color-coded by component. **j.** Projection of voxel responses (**b**) over time for the same trial shown in (**i**), with superimposed outlines of the reconstructed spatial components (W). **k.** Temporal activation patterns (H) of the spatial components (W). **l.** Temporal activation patterns of the voxels defined by the components, as in (**j**)**. m.** Average spatial response centered on the maximally responsive voxel of each spatial component, shown per condition. **n.** Mean spatial component diameter (W) per animal across conditions. **o.** Average temporal response (H) aligned to the respective peak of H for each condition. **p.** Full width at half maximum (FWHM) of the temporal components (H) shown in (**o**), calculated per animal and condition. **q.** Histogram of cross-correlation values between the temporal components (H) and population entropy. All statistical details are provided in Suppl. Table 1.

**Suppl. Fig. 6.**
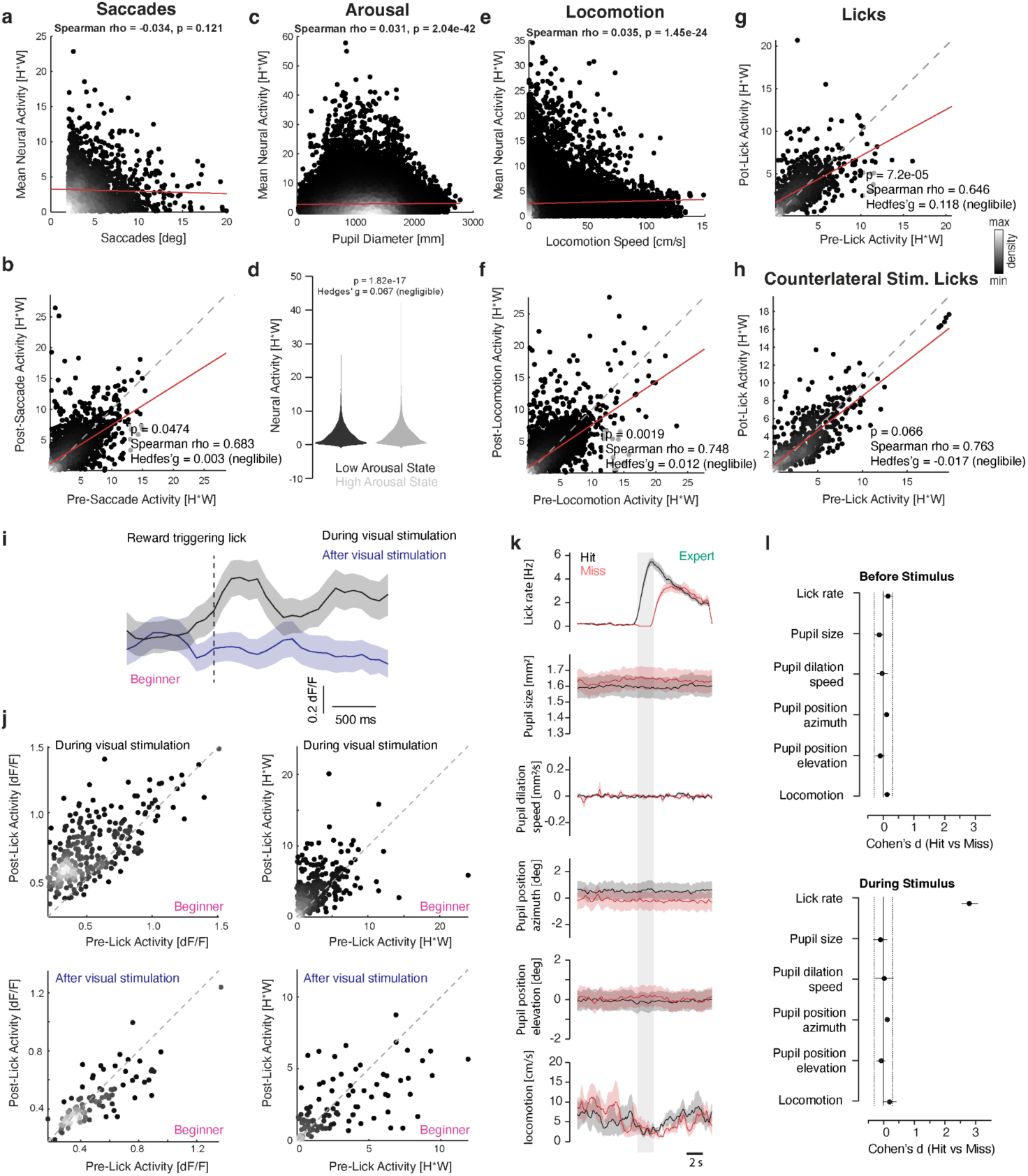
Behavioral, arousal and reward modulation do not explain response changes in reconstructed intrinsic dynamics. **a.** Reconstructed activity changes (H*W) of the imaged area of the superior colliculus are not correlated to the magnitude of saccades, (**b**) saccadic movements (pre- vs post-saccade activity) nor (**c**) arousal (pupil diameter). **d.** The distribution of activities in low and high arousal states (low and high, 10% of the pupil size distribution, respectively) have negligible differences, as (**e**) correlations with locomotion speed, (**f**) the initiation of locomotion (pre- vs post-locomotion activity), (**g**) non-stimulus evoked licking behavior or (**h**) stimulus evoked reactions (stimulus presented contralaterally to the imaged hemisphere to avoid stimulus activity). Red lines: least-squares linear regressions, dashed lines: unity. In **a**, **c**, **e** Spearman ρ and its p-value are reported; in **b**, **d**, **f**, **g**, **h** the p-value, Spearman ρ and Hedges’ g effect size (all negligible) are reported in-panel. Density of observations is indicated by the greyscale color bar (shown next to **g**). **i**. Beginner condition. Top: average ΔF/F of all recorded neurons, aligned to the reward-triggering lick (dashed line) for licks occurring during (black) or after (blue) visual stimulation (mean ± s.e.m.). Note, Beginner animals always get a reward also outside the stimulus window. **j.** Quantification for (**i**) shown for raw ΔF/F (left) and reconstructed intrinsic dynamics (H·W, right). Each dot represents the average response for all recorded neurons per trial. Activity patterns after visual stimulation indicate that licking and reward do not drive intrinsic activity. Activity patterns during visual stimulation show an emerging relationship between intrinsic activity and action, in line with expert responses. **k**. Expert condition. Average behavioral traces aligned to stimulus onset (gray shading) for Hit (black) and Miss (red) trials, including lick rate, pupil size, pupil dilation speed, pupil position (azimuth and elevation), and locomotion speed (mean ± s.e.m.; scale bar, 2 s). **l.** Effect sizes (Cohen’s *d*, Hit vs. Miss) for behavioral variables in (**k**), computed separately before (top) and during (bottom) stimulus presentation. Dotted lines indicate the equivalence bound (|d| = 0.3). Only lick rate during the stimulus shows a large effect, consistent with task design. All statistical details are provided in Suppl. Table 1.

**Suppl. Fig. 7.**
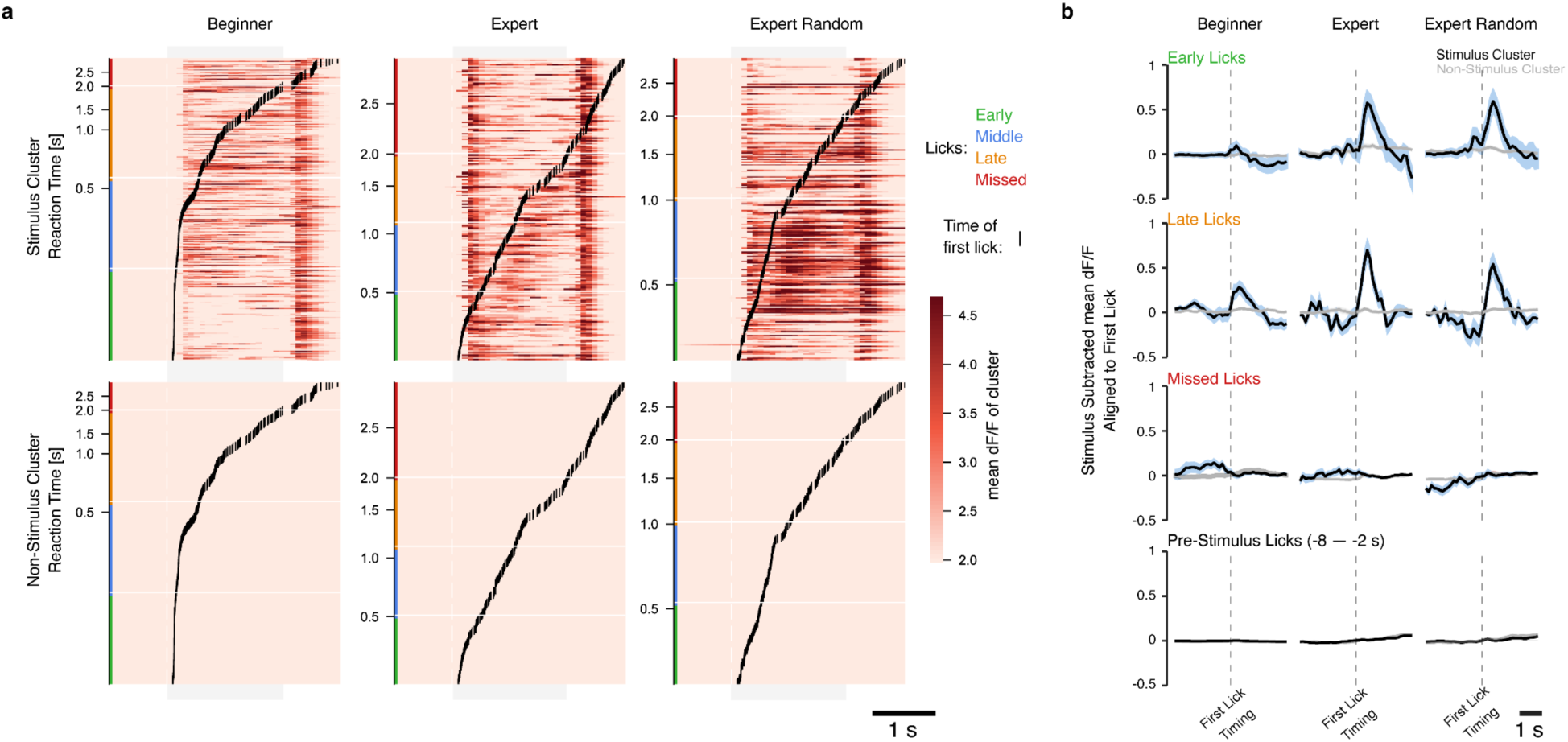
Lick Aligned Blob Activity. **a.** Single-trial activity heatmaps (dF/F) aligned to stimulus onset and sorted by reaction time (time of first lick = black dashes). Trials are color-coded by reaction time tercile: early (green), middle (blue), late (orange) and missed (red). In the Expert conditions, the stimulus cluster visibly exhibits a clear activity increase that aligns with reaction time; a pattern that is absent in the Beginner condition. **b.** Stimulus-subtracted mean dF/F aligned to the time of the first lick separated into distinct lick-timing categories for each condition. Top rows: stimulus cluster activity aligned to licks during the stimulus period exhibits a strong peak near the lick onset in the two expert conditions. Bottom row: pre-stimulus extraneous licks (licks occurring 2-8 s before stimulus onset) show no corresponding activity increase, confirming that the lick-aligned dynamics during the stimulus period reflect task-related blob recruitment rather than a motor artifact of lick activity. All statistical details are provided in Suppl. Table 1.

**Suppl. Fig. 8.**
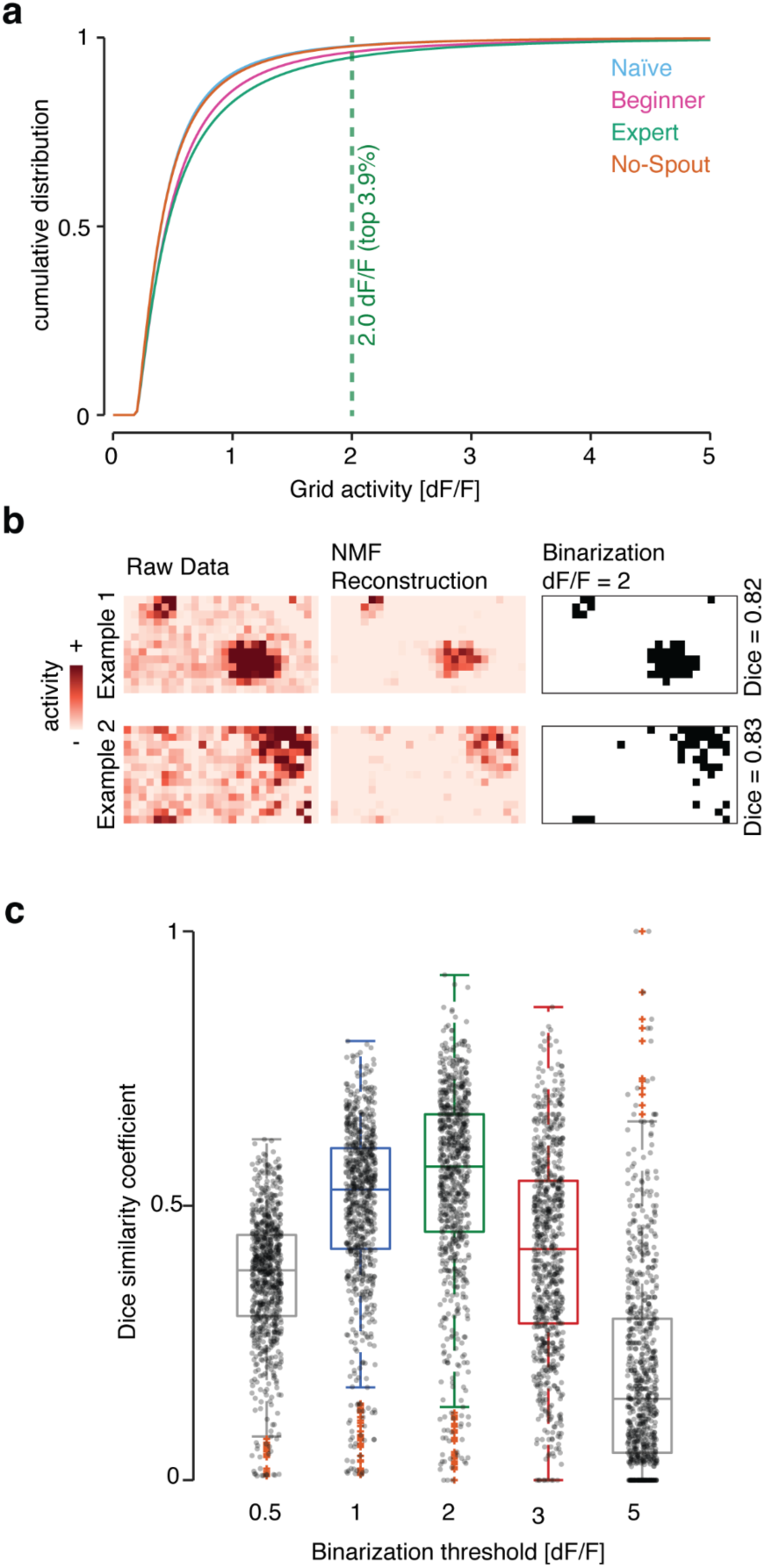
Binarization controls. **a.** Cumulative distribution of voxel activity (dF/F) across all frames and voxels for each condition. The dashed vertical line indicates the binarization threshold of 2.0 dF/F used through the Ising model comparison, which selects the top ∼3.9% of the activity distribution. **b.** Two example trials from an Expert recording show the correspondence between the NMF blob reconstruction (Fig. 3) and the threshold-based binarization. Dice similarity coefficients quantify the spatial overlap of the NMF-detected blob and the pattern after binarization. **c.** Dice similarity coefficient between the NMF reconstruction and the binarized voxel evaluated across five binarization thresholds. The 2.0 dF/F threshold consistently achieves the highest spatial overlap with the NMF reconstruction, which validates it as an appropriate discretization for the Ising model comparison. All statistical details are provided in Suppl. Table 1.

**Suppl. Fig. 9.**
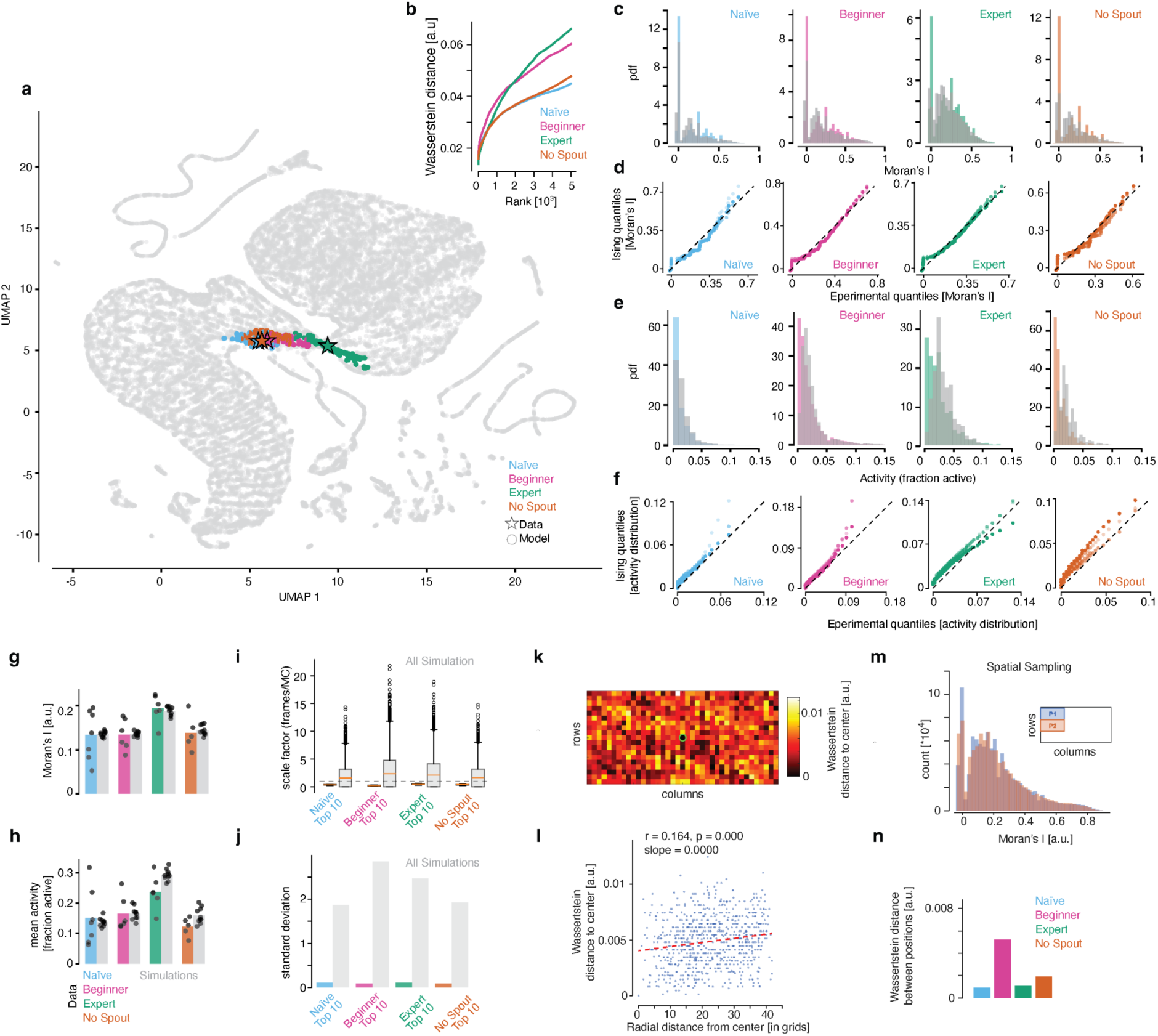
Parameter search and fits. **a.** 2D UMAP embedding of ∼25,000 simulations; colored points mark the top 100 matches per condition, stars denote experimental data. **b.** Wasserstein distance (WD) ranking (top 5,000): only ∼100 simulations achieve low WD before a sharp increase. **c.** Moran’s *I* distributions: experimental data versus best-match simulations. **d.** Quantile-quantile (Q-Q) plots comparing the experimental and top 3 best-match Ising Moran’s I distribution. **e.** Same as (**c**) for the distribution of mean activity (fraction of active voxels per frame). **f.** Same as (**d**) for the activity distributions. **g.** Summary comparison of Moran’s *I* and (**h**) activity for top-10 matches (grey) versus data. **i.** Temporal scale factor (frames per MC sweep) from autocorrelation matching; top-10 versus all simulations. **j.** Variability of scale factors (**i**) per condition; low variance indicates convergence. **k.** Spatial homogeneity test: WD across voxel positions and (**l**) versus radial distance shows no spatial dependence. **m.** Spatial sampling control, exemplified for the Expert condition, confirms homogeneous spatial statistics. Moran’s I distributions computed from two different non-overlapping crop positions (insert). **n.** Mean WD from comparisons as in (**m**) across all conditions. All statistical details are provided in Suppl. Table 1.

**Suppl. Fig. 10.**
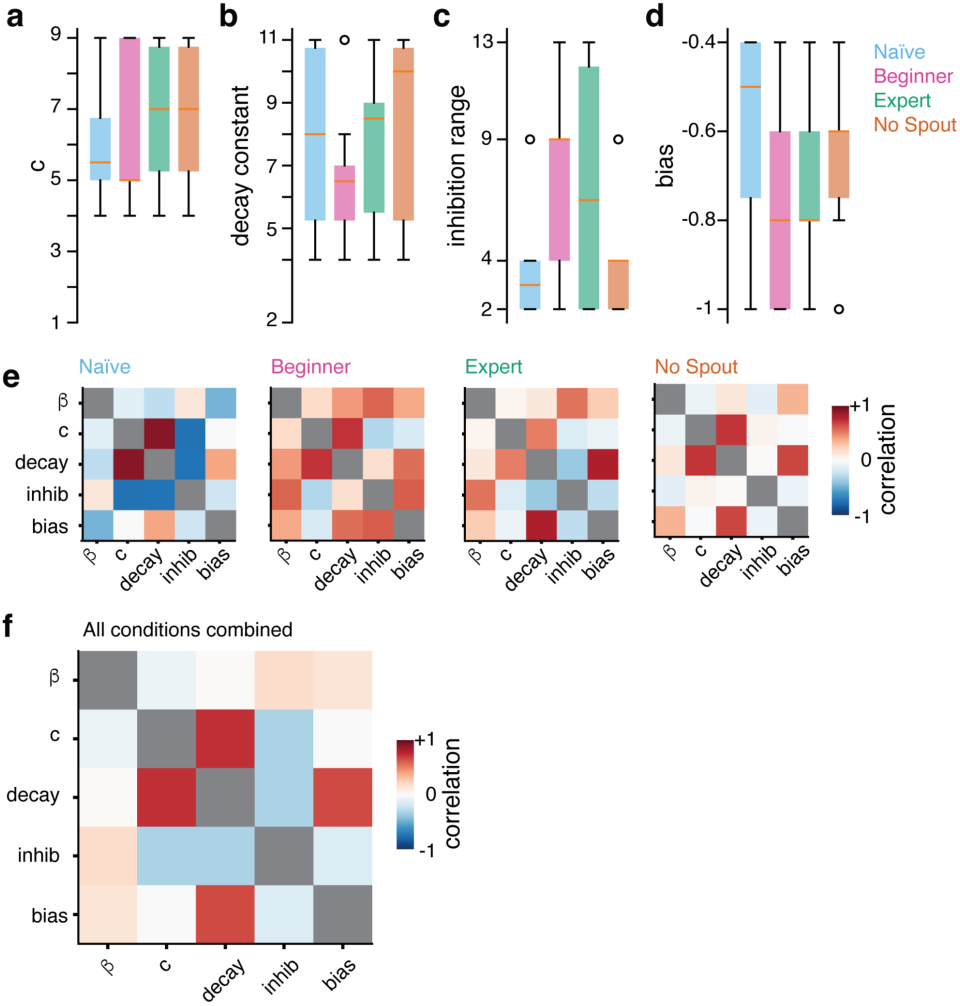
Parameter correlations. **a-d.** Box plots of the four non-β model parameters across the top-ranked simulations for each condition: coupling strength c (**a**), decay constant d (**b**), inhibition range r (**c**) and bias (**d**). Unlike β (Fig. 5f), none of these parameters systematically differ between conditions, indicating that the state-dependent shift in dynamics is not driven by changes in local interaction structure but rather by a global change in excitability. **e.** Pairwise Pearson correlations among parameters within each condition reveal compensatory relationships between non-*β* parameters. **f.** Pooled correlations across conditions (**e**) show consistent compensatory structure independent of *β*, reinforcing *β* as a global control parameter. All statistical details are provided in Suppl. Table 1.

## Supplementary Information

Supplementary Table 1. Statistics Summary

Supplementary Video 1. Example videos for the mean response and three single trials per condition.

Supplementary Video 2. Example videos for the mean response and three single trials per condition (Grid structure, Data same as Suppl. Video 1).

Supplementary Video 3. NMF reconstruction example videos for the mean response and three single trials per condition (Reconstruction of data presented in Suppl. Video 2).

Supplementary Video 4. NMF reconstruction example videos, same as Video 3, but with the mean response subtracted (Reconstruction of data presented in Suppl. Video 2).

Supplementary Video 5. Example (Mouse behavior) Hit/Miss

