## Supplementary material for "Intrinsic Population Dynamics are a Neuronal Substrate for Visual Attention": Statistics Summary

### Figure 1 — Visual Responses Across Learning Stages

#### Panel e — Reaction Times: Predictable vs Random

| Panel | Comparison | Test | p-value | Effect size (Hedges' g) | n |
|---|---|---|---|---|---|
| e | Expert Predictable vs Expert Random | Wilcoxon rank-sum | 0.0155 * | -1.409 | n = 895 vs 565 trials |

#### Panel k — Time-Window Quantification (per-animal)

**Stimulus**

| Panel | Comparison | Test | p-value | Effect size (Hedges' g) | n |
|---|---|---|---|---|---|
| k | All conditions (omnibus) | Kruskal-Wallis | 9.97e-04 *** | N/A | n = 25 observations (animal × condition) |
| k | Naive vs Beginner | Wilcoxon rank-sum | 0.0082 ** | -1.835 | n = 7 vs 6 animals |
| k | Naive vs Expert | Wilcoxon rank-sum | 0.0012 ** | -3.298 | n = 7 vs 6 animals |
| k | Naive vs NoSpout | Wilcoxon rank-sum | 0.4452 n.s. | 0.369 | n = 7 vs 6 animals |
| k | Beginner vs Expert | Wilcoxon rank-sum | 0.2403 n.s. | -0.684 | n = 6 vs 6 animals |
| k | Beginner vs NoSpout | Wilcoxon rank-sum | 0.0152 * | 1.815 | n = 6 vs 6 animals |
| k | Expert vs NoSpout | Wilcoxon rank-sum | 0.0022 ** | 2.921 | n = 6 vs 6 animals |

**StimOnset**

| Panel | Comparison | Test | p-value | Effect size (Hedges' g) | n |
|---|---|---|---|---|---|
| k | All conditions (omnibus) | Kruskal-Wallis | 0.0031 ** | N/A | n = 25 observations (animal × condition) |
| k | Naive vs Beginner | Wilcoxon rank-sum | 0.0734 n.s. | -1.313 | n = 7 vs 6 animals |

| | | | | | |
|---|---|---|---|---|---|
| k | Naive vs Expert | Wilcoxon rank-sum | 0.0012 ** | -3.583 | n = 7 vs 6 animals |
| k | Naive vs NoSpout | Wilcoxon rank-sum | 0.9452 n.s. | 0.108 | n = 7 vs 6 animals |
| k | Beginner vs Expert | Wilcoxon rank-sum | 0.0260 * | -1.596 | n = 6 vs 6 animals |
| k | Beginner vs NoSpout | Wilcoxon rank-sum | 0.1320 n.s. | 1.042 | n = 6 vs 6 animals |
| k | Expert vs NoSpout | Wilcoxon rank-sum | 0.0043 ** | 2.587 | n = 6 vs 6 animals |

**StimOffset**

| Panel | Comparison | Test | p-value | Effect size (Hedges' g) | n |
|---|---|---|---|---|---|
| k | All conditions (omnibus) | Kruskal-Wallis | 6.59e-04 *** | N/A | n = 25 observations (animal × condition) |
| k | Naive vs Beginner | Wilcoxon rank-sum | 0.0047 ** | -2.213 | n = 7 vs 6 animals |
| k | Naive vs Expert | Wilcoxon rank-sum | 0.0012 ** | -3.320 | n = 7 vs 6 animals |
| k | Naive vs NoSpout | Wilcoxon rank-sum | 0.2343 n.s. | 0.893 | n = 7 vs 6 animals |
| k | Beginner vs Expert | Wilcoxon rank-sum | 0.3095 n.s. | -0.614 | n = 6 vs 6 animals |
| k | Beginner vs NoSpout | Wilcoxon rank-sum | 0.0087 ** | 2.296 | n = 6 vs 6 animals |
| k | Expert vs NoSpout | Wilcoxon rank-sum | 0.0022 ** | 3.067 | n = 6 vs 6 animals |

#### Panel l — Exponential Decay Fit

| Panel | Comparison | Test | p-value | Effect size (Hedges' g) | n |
|---|---|---|---|---|---|
| l | All conditions (omnibus) | Kruskal-Wallis | 0.0085 ** | N/A | n = 25 observations (animal × condition) |
| l | Naive vs Beginner | Wilcoxon rank-sum | 0.0221 * | -1.399 | n = 7 vs 6 animals |

| | | | | | |
|---|---|---|---|---|---|
| l | Naive vs Expert | Wilcoxon rank-sum | 0.4452 n.s. | -0.309 | n = 7 vs 6 animals |
| l | Naive vs NoSpout | Wilcoxon rank-sum | 0.1807 n.s. | 1.025 | n = 7 vs 6 animals |
| l | Beginner vs Expert | Wilcoxon rank-sum | 0.0411 * | 1.240 | n = 6 vs 6 animals |
| l | Beginner vs NoSpout | Wilcoxon rank-sum | 0.0087 ** | 2.390 | n = 6 vs 6 animals |
| l | Expert vs NoSpout | Wilcoxon rank-sum | 0.0260 * | 1.568 | n = 6 vs 6 animals |

#### Panel o — CoV per time window across Beginner/Expert/NoSpout

**Pre-stimulus**

| Panel | Comparison | Test | p-value | Effect size (Hedges' g) | n |
|---|---|---|---|---|---|
| o | All conditions (omnibus) | Kruskal-Wallis | 0.0622 n.s. | N/A | n = 18 observations (animal × condition) |
| o | Beginner vs Expert | Wilcoxon rank-sum | 0.0411 * | -1.652 | n = 6 vs 6 animals |
| o | Expert vs NoSpout | Wilcoxon rank-sum | 0.0649 n.s. | 0.911 | n = 6 vs 6 animals |
| o | Beginner vs NoSpout | Wilcoxon rank-sum | 0.8182 n.s. | -0.119 | n = 6 vs 6 animals |

**Stimulus response**

| Panel | Comparison | Test | p-value | Effect size (Hedges' g) | n |
|---|---|---|---|---|---|
| o | All conditions (omnibus) | Kruskal-Wallis | 0.0071 ** | N/A | n = 18 observations (animal × condition) |
| o | Beginner vs Expert | Wilcoxon rank-sum | 0.0087 ** | 1.670 | n = 6 vs 6 animals |
| o | Expert vs NoSpout | Wilcoxon rank-sum | 0.0043 ** | -1.601 | n = 6 vs 6 animals |
| o | Beginner vs NoSpout | Wilcoxon rank-sum | 0.4848 n.s. | -0.363 | n = 6 vs 6 animals |

### Figure 2 — Spatially-Structured Intrinsic Dynamics

#### Panels f-g — Activity Strength vs Distance Correlation

| Panel | Feature | Slope | r |
|---|---|---|---|
| f | Mean activity, strongest 50 cells | -0.04 | -0.739 |
| g | Mean distance between strongest 50 cells | 0.036 | 0.366 |

#### Panel h — Per-Animal Correlation Coefficients (Activity / Distance vs PE)

| Panel | Feature | Test | n (animals) | Median r | p-value (signrank vs 0) |
|---|---|---|---|---|---|
| h | Max activity vs Entropy | Wilcoxon signed-rank vs 0 | n = 6 | -0.232 | 0.0313 * |
| h | Mean distance vs Entropy | Wilcoxon signed-rank vs 0 | n = 6 | 0.162 | 0.0313 * |

#### Panel i — Population Entropy Across Conditions (Trial-Level)

| Panel | Comparison | Test | p-value | Effect size (Hedges' g) | n |
|---|---|---|---|---|---|
| i | Naive vs Beginner (Pre-stimulus Period) | Wilcoxon rank-sum | 2.90e-17 *** | 0.285 | n = 1800 vs 630 trials |
| i | Naive vs Expert (Pre-stimulus Period) | Wilcoxon rank-sum | < 2.22e-308 *** | 0.770 | n = 1800 vs 1176 trials |
| i | Naive vs NoSpout (Pre-stimulus Period) | Wilcoxon rank-sum | 0.3824 n.s. | -0.029 | n = 1800 vs 1050 trials |
| i | Beginner vs Expert (Pre-stimulus Period) | Wilcoxon rank-sum | 1.08e-30 *** | 0.520 | n = 630 vs 1176 trials |
| i | Beginner vs NoSpout (Pre-stimulus Period) | Wilcoxon rank-sum | 1.19e-15 *** | -0.341 | n = 630 vs 1050 trials |

| | | | | | |
|---|---|---|---|---|---|
| i | Expert vs NoSpout (Pre-stimulus Period) | Wilcoxon rank-sum | < 2.22e-308 *** | -0.859 | n = 1176 vs 1050 trials |
| i | Naive vs Expert_P3 (Pre-stimulus Period) | Wilcoxon rank-sum | 3.59e-18 *** | 0.876 | n = 340 vs 137 trials |
| i | Beginner vs Expert_P3 (Pre-stimulus Period) | Wilcoxon rank-sum | 1.15e-05 *** | 0.336 | n = 549 vs 137 trials |
| i | Expert vs Expert_P3 (Pre-stimulus Period) | Wilcoxon rank-sum | 0.0232 * | -0.194 | n = 758 vs 137 trials |
| i | NoSpout vs Expert_P3 (Pre-stimulus Period) | Wilcoxon rank-sum | 6.28e-14 *** | 0.831 | n = 210 vs 137 trials |
| i | Naive vs Beginner (Stimulus period) | Wilcoxon rank-sum | 8.57e-31 *** | 0.480 | n = 1800 vs 630 trials |
| i | Naive vs Expert (Stimulus period) | Wilcoxon rank-sum | < 2.22e-308 *** | 0.655 | n = 1800 vs 1176 trials |
| i | Naive vs NoSpout (Stimulus period) | Wilcoxon rank-sum | 0.7433 n.s. | -0.058 | n = 1800 vs 1050 trials |
| i | Beginner vs Expert (Stimulus period) | Wilcoxon rank-sum | 0.0130 * | 0.155 | n = 630 vs 1176 trials |
| i | Beginner vs NoSpout (Stimulus period) | Wilcoxon rank-sum | 5.32e-30 *** | -0.568 | n = 630 vs 1050 trials |
| i | Expert vs NoSpout (Stimulus period) | Wilcoxon rank-sum | < 2.22e-308 *** | -0.764 | n = 1176 vs 1050 trials |

##### Panel j — Fold-Change in Population Entropy to Naive (Per-Animal)

**Pairwise Wilcoxon rank-sum:**

| Panel | Comparison | Test | n | p-value | Significance |
|---|---|---|---|---|---|
| j | Beginner_P1 vs Expert_P1 | Wilcoxon rank-sum | n = 7 vs 5 | 0.0025 ** | ** |
| j | Beginner_P1 vs Expert_P3 | Wilcoxon rank-sum | n = 7 vs 5 | 0.0101 * | * |
| j | Beginner_P1 vs NoSpout_P1 | Wilcoxon rank-sum | n = 7 vs 7 | 0.5350 n.s. | n.s. |
| j | Expert_P1 vs Expert_P3 | Wilcoxon rank-sum | n = 5 vs 5 | 0.3095 n.s. | n.s. |
| j | Expert_P1 vs NoSpout_P1 | Wilcoxon rank-sum | n = 5 vs 7 | 0.0025 ** | ** |
| j | Expert_P3 vs NoSpout_P1 | Wilcoxon rank-sum | n = 5 vs 7 | 0.0051 ** | ** |

#### Panel k — SVM Classification Accuracy (5-fold CV)

| Panel | Comparison | Test | n | Accuracy | Chance |
|---|---|---|---|---|---|
| k | Naive vs Expert (2-class) | Kernelized SVM, 5-fold CV | n = 2976 trials | 86.9% | 50% |
| k | Beginner vs Expert (2-class) | Kernelized SVM, 5-fold CV | n = 1806 trials | 87.3% | 50% |
| k | NoSpout vs Expert (2-class) | Kernelized SVM, 5-fold CV | n = 2226 trials | 80.6% | 50% |
| k | All conditions (4-class) | Kernelized SVM, 5-fold CV | n = 4656 trials | 65.3% | 25% |

#### Panel m — FWHM of Spatial Response Across Conditions

| Panel | Mode | Comparison | Test | p-value | Effect size (Hedges' g) | n |
|---|---|---|---|---|---|---|
| m | EntropyTriggered | Across conditions (omnibus) | Kruskal-Wallis | 0.5811 n.s. | — | — |
| m | EntropyTriggered | Naive vs Beginner | Wilcoxon rank-sum | 0.2976 n.s. | -0.504 | n = 12 vs 8 |
| m | EntropyTriggered | Naive vs Expert | Wilcoxon rank-sum | 0.6650 n.s. | -0.025 | n = 12 vs 12 |
| m | EntropyTriggered | Naive vs NoSpout | Wilcoxon rank-sum | 0.5358 n.s. | -0.654 | n = 12 vs 7 |

| | | | | | | |
|---|---|---|---|---|---|---|
| m | EntropyTriggered | Beginner vs Expert | Wilcoxon rank-sum | 0.2976 n.s. | 0.367 | n = 8 vs 12 |
| m | EntropyTriggered | Beginner vs NoSpout | Wilcoxon rank-sum | 0.8665 n.s. | -0.249 | n = 8 vs 7 |
| m | EntropyTriggered | Expert vs NoSpout | Wilcoxon rank-sum | 0.3845 n.s. | -0.545 | n = 12 vs 7 |
| m | StimPeriod | Across conditions (omnibus) | Kruskal-Wallis | 0.0060 ** | — | — |
| m | StimPeriod | Naive vs Beginner | Wilcoxon rank-sum | 0.0186 * | -1.619 | n = 12 vs 8 |
| m | StimPeriod | Naive vs Expert | Wilcoxon rank-sum | 0.0120 * | -1.188 | n = 12 vs 12 |
| m | StimPeriod | Naive vs NoSpout | Wilcoxon rank-sum | 0.5358 n.s. | 0.243 | n = 12 vs 7 |
| m | StimPeriod | Beginner vs Expert | Wilcoxon rank-sum | 0.1535 n.s. | 0.814 | n = 8 vs 12 |
| m | StimPeriod | Beginner vs NoSpout | Wilcoxon rank-sum | 0.0205 * | 1.414 | n = 8 vs 7 |
| m | StimPeriod | Expert vs NoSpout | Wilcoxon rank-sum | 0.0283 * | 1.148 | n = 12 vs 7 |

##### Panel o — FWHM of Spatial Response (Stimulus-Triggered)

| Panel | Comparison | Test | p-value | Effect size (Hedges' g) | n |
|---|---|---|---|---|---|
| o | Across conditions (omnibus) | Kruskal-Wallis | 0.5828 n.s. | — | — |
| o | Naive vs Beginner | Wilcoxon rank-sum | 0.2631 n.s. | -0.529 | n = 12 vs 8 |
| o | Naive vs Expert | Wilcoxon rank-sum | 0.7508 n.s. | -0.094 | n = 12 vs 12 |
| o | Naive vs NoSpout | Wilcoxon rank-sum | 0.2268 n.s. | -0.464 | n = 12 vs 7 |
| o | Beginner vs Expert | Wilcoxon rank-sum | 0.5627 n.s. | 0.471 | n = 8 vs 12 |
| o | Beginner vs NoSpout | Wilcoxon rank-sum | 0.5159 n.s. | 0.047 | n = 8 vs 7 |

| | | | | | |
|---|---|---|---|---|---|
| o | Expert vs NoSpout | Wilcoxon rank-sum | 0.6064 n.s. | -0.405 | n = 12 vs 7 |

### Figure 3 — Blob-Like Activity During Stimulation

#### Panel c — Jaccard Fidelity: Blobs vs Visual Response

| Panel | Comparison | Test | p-value | Effect size (Hedges' g) | n |
|---|---|---|---|---|---|
| c | Mean Jaccard: Blobs vs Visual Response | Wilcoxon rank-sum | 3.66e-05 *** | -5.842 | n = 12 vs 12 recordings |

#### Panel e — Event Density: Near vs Far vs Contralateral (Expert)

| Panel | Period | Comparison | Test | p-value | Effect size (Hedges' g) | n |
|---|---|---|---|---|---|---|
| e | Before Stimulus | Near vs Far | Wilcoxon rank-sum | 1.0000 n.s. | 0.174 | n = 12 vs 12 |
| e | Before Stimulus | Near vs Contralateral | Wilcoxon rank-sum | 0.7871 n.s. | 0.046 | n = 12 vs 8 |
| e | Before Stimulus | Far vs Contralateral | Wilcoxon rank-sum | 0.9079 n.s. | -0.148 | n = 12 vs 8 |
| e | During Stimulus | Near vs Far | Wilcoxon rank-sum | 7.31e-04 *** | 1.724 | n = 12 vs 12 |
| e | During Stimulus | Near vs Contralateral | Wilcoxon rank-sum | 0.0062 ** | 1.502 | n = 12 vs 8 |
| e | During Stimulus | Far vs Contralateral | Wilcoxon rank-sum | 0.5628 n.s. | -0.318 | n = 12 vs 8 |

#### Panel g — Event Density: Near vs Far vs Contralateral (No Spout)

| Panel | Period | Comparison | Test | p-value | Effect size (Hedges' g) | n |
|---|---|---|---|---|---|---|
| g | Before Stimulus | Near vs Far | Wilcoxon rank-sum | 0.6200 n.s. | -0.386 | n = 7 vs 7 |
| g | Before Stimulus | Near vs Contralateral | Wilcoxon rank-sum | 0.3176 n.s. | -0.594 | n = 7 vs 7 |
| g | Before Stimulus | Far vs Contralateral | Wilcoxon rank-sum | 0.7104 n.s. | -0.193 | n = 7 vs 7 |
| g | During Stimulus | Near vs Far | Wilcoxon rank-sum | 0.3939 n.s. | 0.797 | n = 6 vs 6 |
| g | During Stimulus | Near vs Contralateral | Wilcoxon rank-sum | 0.6991 n.s. | 0.836 | n = 6 vs 6 |

| | | | | | | |
|---|---|---|---|---|---|---|
| g | During Stimulus | Far vs Contralateral | Wilcoxon rank-sum | 0.6991 n.s. | 0.167 | n = 6 vs 6 |

### Figure 4 — Hit/Miss and Behavioral Performance

#### Panel c — Hit vs Miss dF/F (per-cell, Expert)

| Panel | Comparison | Test | p-value | Effect size (Hedges' g) | n |
|---|---|---|---|---|---|
| c | Activity: Hit vs Miss (Expert, peak frame, all cells) | Wilcoxon signed-rank | < 2.22e-308 *** | 0.440 | n = 1618 cells |

#### Panel d — Hit vs Miss dF/F (per-animal, Expert) + mean modulation

| Panel | Comparison | Test | p-value | Effect size (Hedges' g) | n | Mean modulation |
|---|---|---|---|---|---|---|
| d | Activity: Hit vs Miss (Expert, peak frame, per recording) | Wilcoxon signed-rank | 0.0078 ** | 1.404 | n = 8 recordings | 34.91% |

#### Panel e — Hit vs Miss SVM Classifier (Pre-stimulus PE)

| Panel | RT cutoff | Test | Sampling | Accuracy | Chance |
|---|---|---|---|---|---|
| e | 2 s | Kernelized SVM, 5-fold CV | class balanced | 64% | 50% |
| e | 3 s | Kernelized SVM, 5-fold CV | class balanced | 69% | 50% |
| e | 4 s | Kernelized SVM, 5-fold CV | class balanced | 73% | 50% |
| e | 5 s | Kernelized SVM, 5-fold CV | class balanced | 87% | 50% |

#### Panel g — Modulation Index (Center – Onset), Hit vs Hit Random

| Panel | Comparison | Test | p-value | Effect size (Hedges' g) | n |
|---|---|---|---|---|---|
| g | ModIdx (Center – Onset): Hit (Expert) vs Hit Random (ExpertRandom) | Wilcoxon rank-sum | < 2.22e-308 *** | -0.523 | n = 1618 vs 1975 cells |

#### Panel h — Peak timing vs reaction time (StimCluster cells)

| Panel | Condition | Test | n | r | p-value |
|---|---|---|---|---|---|
| h | Overall | Pearson correlation | n = 626 | 0.280 | 9.89e-13 *** |
| h | Beginner | Pearson correlation | n = 288 | 0.126 | 0.0319 * |
| h | Expert | Pearson correlation | n = 151 | 0.346 | 1.37e-05 *** |
| h | ExpertRandom | Pearson correlation | n = 187 | 0.452 | 8.33e-11 *** |

#### Panel j — Stimulus Cluster Size, Small-Stim Hit vs Miss + NoSpout

| Panel | Condition / Comparison | Test | p-value | Effect size (Hedges' g) | n |
|---|---|---|---|---|---|
| j | SmallStimBeginner: Hit vs Miss | Wilcoxon rank-sum | 0.8955 n.s. | -0.038 | n = 182 vs 78 trials |
| j | SmallStimExpert: Hit vs Miss | Wilcoxon rank-sum | 1.95e-07 *** | 0.642 | n = 109 vs 152 trials |
| j | Beginner Hit vs Expert Hit | Wilcoxon rank-sum | 5.39e-07 *** | -0.759 | n = 182 vs 109 trials |
| j | Beginner Miss vs Expert Hit | Wilcoxon rank-sum | 2.49e-06 *** | -0.689 | n = 78 vs 108 trials |
| j | Expert (Hit) vs NoSpout (pool) | Wilcoxon rank-sum | 2.16e-07 *** | 0.646 | n = 109 vs 539 trials |
| j | Beginner (Hit) vs NoSpout (pool) | Wilcoxon rank-sum | 0.2363 n.s. | -0.142 | n = 186 vs 539 trials |

#### Panel l — Event density Near / Far, Hit vs Miss (peak frame)

| Panel | Region | Comparison | Test | p-value | Effect size (Hedges' g) | n |
|---|---|---|---|---|---|---|
| l | Near | Hit vs Miss | Wilcoxon signed-rank | 0.0156 * | 1.78 | n = 7 recordings |
| l | Far | Hit vs Miss | Wilcoxon signed-rank | 0.0625 n.s. | 0.87 | n = 7 recordings |

#### Panel m — Peak Activity Timing vs Reaction Time (Small Stimulus)

| Panel | Population | Test | n | r | $R^2$ | p-value |
|---|---|---|---|---|---|---|

| | | | | | | |
|---|---|---|---|---|---|---|
| m | Visually-responsive cluster (StimCluster) | Pearson correlation | n = 123 trials | -0.026 | 0.001 | 0.774 n.s. |
| m | Non-stimulus cluster (NonStimCluster) | Pearson correlation | n = 123 trials | 0.524 | 0.275 | 4.77e-10 *** |

### Figure 5 — Ising Model

#### Model Description

- 2D Ising model with local excitatory-inhibitory interactions on a 39 x 78 lattice (3x experimental grid dimensions) with periodic boundary conditions
- Heat-bath algorithm for spin dynamics
- 24,624 parameter combinations explored (beta: 19 values [0.4-0.8], c: [1,2,3,4,5,6,7,8,9], decay constant: [2,4,5,6,7,8,9,10,11], r: [2,4,9,13], b: [-1.0,-0.8,-0.6,-0.4])
- Each simulation: ≥2,000 burn-in (adaptive: max(2000, 7τ)) + 100,000 recording Monte Carlo sweeps
- Model-data matching via two metrics: Moran's I (spatial autocorrelation) and mean activity
- Combined matching used equal weights for these metrics

---

##### Panel d — Model↔Data composite matching score

Descriptive composite Wasserstein score (Moran's I + activity distribution, equal weights) between each condition's data and its best-matching Ising simulation. Smaller = better match; values are rank-normalised across the 24,527-combo parameter grid, so negative W means "better than the grid median".

| Panel | Comparison | Test | n | Wasserstein distance (top-1) | Wasserstein distance (top-10 mean) |
|---|---|---|---|---|---|
| d | Model↔Data match score (Naive) | Composite Wasserstein | 24,527 param combos | W = -1.121 | W = -1.090 |
| d | Model↔Data match score (Beginner) | Composite Wasserstein | 24,527 | W = -1.379 | W = -1.343 |
| d | Model↔Data match score (Expert) | Composite Wasserstein | 24,527 | W = -1.544 | W = -1.524 |
| d | Model↔Data match score (NoSpout) | Composite Wasserstein | 24,527 | W = -1.140 | W = -1.132 |

##### Panel f — Inverse temperature β across conditions

Pairwise Wilcoxon rank-sum (Mann-Whitney U) on the 10 top-match parameter sets per condition; Kruskal–Wallis omnibus over all four conditions. Significance reflects the raw p-value.

| Panel | Comparison | Test | n | U | p-value | Significance |
|---|---|---|---|---|---|---|
| f | β across Naive/Beginner/Expert/NoSpout | Kruskal-Wallis | 40 (10/cond) | H = 27.34 | 5.00e-06 *** | *** |
| f | β: Naive vs Beginner | Mann-Whitney U | 10 vs 10 | 86 | 7.15e-03 ** | ** |

| | | | | | | |
|---|---|---|---|---|---|---|
| f | β: Naive vs Expert | Mann-Whitney U | 10 vs 10 | 100 | 1.56e-04 *** | *** |
| f | β: Naive vs NoSpout | Mann-Whitney U | 10 vs 10 | 78 | 2.79e-02 * | * |
| f | β: Beginner vs Expert | Mann-Whitney U | 10 vs 10 | 100 | 1.89e-04 *** | *** |
| f | β: Beginner vs NoSpout | Mann-Whitney U | 10 vs 10 | 38 | 3.81e-01 n.s. | n.s. |
| f | β: Expert vs NoSpout | Mann-Whitney U | 10 vs 10 | 0 | 1.37e-04 *** | *** |

### Figure 6 — State-Dependent Amplification (Ising perturbations)

#### Panel d — Δ Fraction Active: Expert vs No-Spout

| Panel | Type | Comparison | Test | p-value | Effect size (Hedges' g) | n |
|---|---|---|---|---|---|---|
| d | Data | Δ Fraction Active: Expert vs NoSpout | Wilcoxon rank-sum | 0.00932 ** | 1.457 | n = 8 vs 7 rec |
| d | Model | Δ Fraction Active: Expert vs NoSpout | Wilcoxon rank-sum | 1.60e-16 *** | 1.595 | n = 50 vs 50 sim |

#### Panel e — Sensitivity Threshold (EC50 from Hill fits, Ising Gating)

| Panel | Condition | Sensitivity Threshold (voxels²) | Hill n | Fit R² |
|---|---|---|---|---|
| e | Expert | 12.519 | 1.001 | 0.683 |
| e | NoSpout | 39.909 | 2.168 | 0.798 |

| Panel | Comparison | Test | p-value | Effect size (Hedges' g) | n |
|---|---|---|---|---|---|
| e | Expert vs NoSpout | Wilcoxon rank-sum | 1.83e-04 *** | -3.845 | n = 10 vs 10 sim |

#### Panel g — Max Fraction Active (small stim): Hit / Miss / NoSpout

| Panel | Type | Comparison | Test | p-value | Effect size (Hedges' g) | n |
|---|---|---|---|---|---|---|
| g | Data | Hit vs Miss | Wilcoxon rank-sum | 4.95e-09 *** | 0.497 | n = 118 vs 170 trial |
| g | Data | Hit vs NoSpout | Wilcoxon rank-sum | 1.47e-15 *** | 0.801 | n = 118 vs 600 trial |
| g | Data | Miss vs NoSpout | Wilcoxon rank-sum | 0.2074 n.s. | 0.191 | n = 170 vs 600 trial |
| g | Model | Expert vs NoSpout | Wilcoxon rank-sum | 1.71e-05 *** | 0.165 | n = 1000 vs 1000 rep |

### Extended Data Figure 1 — Attentional Focus Shifts (Cueing & Repeated Stimulation)

#### Panel b — Reaction Time After Location Switch

| Panel | Comparison | Test | p-value | Effect size (Hedges' g) | n |
|---|---|---|---|---|---|
| b | RT: Trial 1 vs Trial 2 (after switch) | Wilcoxon rank-sum | 0.0085 ** | 0.406 | n = 60 vs 57 trials |
| b | RT: Trial 1 vs Trial 3 (after switch) | Wilcoxon rank-sum | 0.0049 ** | 0.471 | n = 60 vs 60 trials |
| b | RT: Trial 1 vs Trials 4+ (after switch) | Wilcoxon rank-sum | 5.49e-06 *** | 0.460 | n = 60 vs 406 trials |

#### Panel d — Reaction Time: Expert (uncued) / Expert Cued / Expert Wrong Cue

| Panel | Comparison | Test | p-value | Effect size (Hedges' g) | n |
|---|---|---|---|---|---|
| d | RT: Expert (uncued) vs Expert Cued | Wilcoxon rank-sum | 7.49e-22 *** | +0.099 | n = 889 vs 914 trials |
| d | RT: Expert (uncued) vs Expert Wrong Cue | Wilcoxon rank-sum | 0.583 (n.s.) | –0.303 | n = 889 vs 22 trials |
| d | RT: Expert Cued vs Expert Wrong Cue | Wilcoxon rank-sum | 0.029 * | –0.343 | n = 914 vs 22 trials |

#### Panel h — Reaction Time at Imaging Retinotopy: Expert / Random / Small

| Panel | Comparison | Test | p-value | Effect size (Hedges' g) | n |
|---|---|---|---|---|---|
| h | RT: Expert vs Expert Random | Wilcoxon rank-sum | 8.64e-22 *** | –0.682 | n = 9 116 vs 268 trials |
| h | RT: Expert vs Expert Small (2°) | Wilcoxon rank-sum | 4.61e-162 *** | –0.564 | n = 9 116 vs 1 910 trials |
| h | RT: Expert Random vs Expert Small | Wilcoxon rank-sum | 0.170 (n.s.) | +0.167 | n = 268 vs 1 910 trials |

#### Panel j — Hit Rate: Expert / Random / Small

| Panel | Comparison | Test | p-value | Effect size (Hedges' g) | n |
|---|---|---|---|---|---|
| j | Hit rate: Expert vs Expert Random | Wilcoxon rank-sum | 6.61e-06 *** | 1.001 | n = 118 vs 33 sessions |
| j | Hit rate: Expert vs Expert Small | Wilcoxon rank-sum | 9.71e-11 *** | 1.679 | n = 118 vs 33 sessions |
| j | Hit rate: Expert Random vs Expert Small | Wilcoxon rank-sum | 0.0008 *** | 1.098 | n = 33 vs 33 sessions |

### Extended Data Figure 2 — Behavioural, Arousal and Reward Modulation Controls

#### Panels a–b — Saccades vs Neural Activity

| Panel | Comparison | Test | n | p-value | Spearman rho | Hedges' g |
|---|---|---|---|---|---|---|
| a | Saccade (pupil movement) vs Mean Activity (dF/F) | Spearman correlation | n = 15019 movement frames | 0.443 n.s. | –0.006 | — |
| b | Pre vs Post Saccade Activity | Wilcoxon signed-rank | n = 1792 events | < 2.22e-308 *** | 0.67 | 0.03 |

#### Panels c–d — Arousal (Pupil Size) vs Neural Activity

| Panel | Comparison | Test | n | p-value | Spearman rho | Hedges' g |
|---|---|---|---|---|---|---|
| c | Pupil Diameter vs Mean Activity (dF/F) | Spearman correlation | n = 303 206 frames | < 2.22e-308 *** | 0.064 | — |
| d | Low vs High Arousal State Activity | Wilcoxon rank-sum | High n = 87 580, Low n = 73 924 frames | < 2.22e-308 *** | — | 0.167 |

#### Panels e–f — Locomotion vs Neural Activity

| Panel | Comparison | Test | n | p-value | Spearman rho | Hedges' g |
|---|---|---|---|---|---|---|
| e | Locomotion Speed vs Mean Activity (dF/F) | Spearman correlation | n = 9132 frames | < 2.22e-308 *** | 0.194 | — |
| f | Pre vs Post Locomotion Activity | Wilcoxon signed-rank | n = 62994 events | < 2.22e-308 *** | 0.877 | 0.054 |

#### Panels g–h — Licking vs Neural Activity

| Panel | Comparison | Test | n | p-value | Spearman rho | Hedges' g |
|---|---|---|---|---|---|---|
| g | Pre vs Post Lick Activity (non-stimulus / extraneous licks) | Wilcoxon signed-rank | n = 1634 events | < 2.22e-308 *** | 0.64 | 0.093 |
| h | Pre vs Post Lick Activity (contralateral stimulus licks) | Wilcoxon signed-rank | n = 125 events | < 2.22e-308 *** | 0.675 | 0.315 |

### Extended Data Figure 4 — Behavioural / Arousal Controls for Population Entropy

#### Panels a–b — Saccades vs Population Entropy

| Panel | Comparison | Test | n | p-value | Spearman rho | Hedges' g |
|---|---|---|---|---|---|---|
| a | Saccade (pupil movement) vs Population Entropy | Spearman correlation | n = 8518 movement frames | 0.00149 ** | –0.031 | — |
| b | Pre vs Post Saccade Entropy | Wilcoxon signed-rank | n = 1791 events | 0.00288 ** | 0.742 | 0.002 |

#### Panels c–d — Arousal (Pupil Size) vs Population Entropy

| Panel | Comparison | Test | n | p-value | Spearman rho | Hedges' g |
|---|---|---|---|---|---|---|
| c | Pupil Diameter vs Population Entropy | Spearman correlation | n = 300 440 frames | < 2.22e-308 *** | –0.128 | — |
| d | Low vs High Arousal State Entropy | Wilcoxon rank-sum | High n = 86 022, Low n = 73 675 frames | < 2.22e-308 *** | — | –0.321 |

#### Panels e–f — Locomotion vs Population Entropy

| Panel | Comparison | Test | n | p-value | Spearman rho | Hedges' g |
|---|---|---|---|---|---|---|
| e | Locomotion Speed vs Population Entropy | Spearman correlation | n = 9119 frames | 1.08e-31 *** | –0.122 | — |
| f | Pre vs Post Locomotion Entropy | Wilcoxon signed-rank | n = 7157 events | 7.4e-05 *** | 0.861 | 0.033 |

#### Panels g–h — Licking vs Population Entropy

| Panel | Comparison | Test | n | p-value | Spearman rho | Hedges' g |
|---|---|---|---|---|---|---|

| | | | | | | |
|---|---|---|---|---|---|---|
| g | Pre vs Post Lick Entropy (non-stimulus / extraneous licks) | Wilcoxon signed-rank | n = 1634 events | 1.02e-12 *** | 0.725 | 0.139 |
| h | Pre vs Post Lick Entropy (contralateral stimulus licks) | Wilcoxon signed-rank | n = 102 events | 7.06e-10 *** | 0.764 | 0.463 |

### Extended Data Figure 5 — NMF Decomposition Validation

#### Panel d — Pearson Correlation Distributions (Within vs Across Voxels)

Per-grid-cell mean pairwise Pearson correlation between cell trace pairs. **Actual** = pairs of cells within the same voxel (Grid40, 40 × 40 µm). **Random** = same-size groups drawn from random cells across the imaging field. Shown for both All-Cells and the visually responsive Stim-Cells subset, for stimuli at Position 1 (P1) and Position 3 (P3).

| Panel | Position | Group | Comparison | Test | n (Actual + Random) | p-value | Hedges' g |
|---|---|---|---|---|---|---|---|
| d | P1 | All cells | Actual (within voxel) vs Random (across voxels) | Wilcoxon rank-sum | 13 714 voxels | < 2.22e-308 *** | +2.534 |
| d | P1 | Stim cells | Actual (within voxel) vs Random (across voxels) | Wilcoxon rank-sum | 2 556 voxels | < 2.22e-308 *** | +3.005 |
| d | P3 | All cells | Actual (within voxel) vs Random (across voxels) | Wilcoxon rank-sum | 10 574 voxels | < 2.22e-308 *** | +2.427 |
| d | P3 | Stim cells | Actual (within voxel) vs Random (across voxels) | Wilcoxon rank-sum | 1 890 voxels | < 2.22e-308 *** | +2.898 |

#### Panel n — Mean Spatial Component (W) Diameter Across Conditions

Per-recording mean component diameter, one value per recording. Test is Kruskal-Wallis omnibus across the four conditions, followed by 6 Wilcoxon rank-sum pairwise comparisons with Holm-Bonferroni adjustment over those 6 tests. Hedges' g sign is `(group_a − group_b) / pooled_sd`.

| Panel | Comparison | Test | n | p-value | Hedges' g | Significance |
|---|---|---|---|---|---|---|
| n | W diameter across Naive/Beginner/Expert/NoSpout | Kruskal-Wallis | 56 (per recording) | 0.116 n.s. | — | n.s. |
| n | Naive vs Beginner | Wilcoxon rank-sum | 25 | 0.270 n.s. | −0.297 | n.s. |

| | | | | | | |
|---|---|---|---|---|---|---|
| n | Naive vs Expert | Wilcoxon rank-sum | 33 | 0.050 n.s. | –0.611 | n.s. (raw *) |
| n | Naive vs NoSpout | Wilcoxon rank-sum | 30 | 0.048 * | –0.861 | n.s. (raw *) |
| n | Beginner vs Expert | Wilcoxon rank-sum | 26 | 0.590 n.s. | –0.350 | n.s. |
| n | Beginner vs NoSpout | Wilcoxon rank-sum | 23 | 0.270 n.s. | –0.640 | n.s. |
| n | Expert vs NoSpout | Wilcoxon rank-sum | 31 | 0.565 n.s. | –0.455 | n.s. |

##### Panel p — Temporal H Component FWHM Across Conditions

Per-recording mean H-component FWHM. Same test family and sign convention as panel n.

| Panel | Comparison | Test | n | p-value | Hedges' g | Significance |
|---|---|---|---|---|---|---|
| p | H FWHM across Naive/Beginner/Expert/NoSpout | Kruskal-Wallis | 56 (per recording) | 0.261 n.s. | — | n.s. |
| p | Naive vs Beginner | Wilcoxon rank-sum | 25 | 0.713 n.s. | –0.284 | n.s. |
| p | Naive vs Expert | Wilcoxon rank-sum | 33 | 0.228 n.s. | +0.270 | n.s. |
| p | Naive vs NoSpout | Wilcoxon rank-sum | 30 | 0.190 n.s. | +0.463 | n.s. |
| p | Beginner vs Expert | Wilcoxon rank-sum | 26 | 0.106 n.s. | +0.446 | n.s. |
| p | Beginner vs NoSpout | Wilcoxon rank-sum | 23 | 0.197 n.s. | +0.726 | n.s. |
| p | Expert vs NoSpout | Wilcoxon rank-sum | 31 | 0.620 n.s. | +0.054 | n.s. |

### Extended Data Figure 6 — Behavioural Controls for Reconstructed Intrinsic Dynamics (H·W)

#### Panels a–b — Saccades vs Reconstructed Activity (H·W)

| Panel | Comparison | Test | n | p-value | Hedges' g | Spearman rho |
|---|---|---|---|---|---|---|
| a | Saccade (pupil movement) vs Mean H·W Activity | Spearman correlation | — | 0.121 n.s. | — | –0.034 |
| b | Pre vs Post Saccade H·W Activity | Spearman correlation (pre vs post) | — | 0.0474 * | 0.003 | 0.683 |

#### Panels c–d — Arousal (Pupil Size) vs Reconstructed Activity (H·W)

| Panel | Comparison | Test | n | p-value | Hedges' g | Spearman rho |
|---|---|---|---|---|---|---|
| c | Pupil Diameter vs Mean H·W Activity | Spearman correlation | — | 2.04e-42 *** | — | 0.031 |
| d | Low vs High Arousal State H·W Activity | Wilcoxon rank-sum | — | 1.82e-17 *** | 0.067 | — |

#### Panels e–f — Locomotion vs Reconstructed Activity (H·W)

| Panel | Comparison | Test | n | p-value | Hedges' g | Spearman rho |
|---|---|---|---|---|---|---|
| e | Locomotion Speed vs Mean H·W Activity | Spearman correlation | — | 1.45e-24 *** | — | 0.035 |
| f | Pre vs Post Locomotion H·W Activity | Spearman correlation (pre vs post) | — | 1.92e-04 *** | 0.012 | 0.748 |

#### Panels g–h — Licking vs Reconstructed Activity (H·W)

| Panel | Comparison | Test | n | p-value | Hedges' g | Spearman rho |
|---|---|---|---|---|---|---|

| | | | | | | |
|---|---|---|---|---|---|---|
| g | Pre vs Post Lick H·W Activity (ipsilateral stim licks) | Spearman correlation (pre vs post) | — | 7.23e-05 *** | 0.118 | 0.646 |
| h | Pre vs Post Lick H·W Activity (counterlateral stim licks) | Spearman correlation (pre vs post) | — | 0.066 n.s. | –0.017 | 0.763 |

#### Panel j — Beginner Pre-Lick vs Post-Lick Activity (Lick-Triggered)

| Panel | Period | Metric | Test | n | p-value | Hedges' g | Spearman rho |
|---|---|---|---|---|---|---|---|
| j | During visual stimulation (Hit licks) | Raw dF/F | Spearman + Wilcoxon signed-rank | 288 trials | < 2.22e-308 (rho); signrank p = 8.55e-34 *** | 0.750 | 0.613 |
| j | During visual stimulation (Hit licks) | Reconstructed H·W | Spearman + Wilcoxon signed-rank | 279 trials | < 2.22e-308 (rho); signrank p = 2.72e-13 *** | 0.384 | 0.538 |
| j | After visual stimulation (Miss licks) | Raw dF/F | Spearman + Wilcoxon signed-rank | 105 trials | < 2.22e-308 (rho); signrank p = 3.22e-03 *** | –0.203 | 0.829 |
| j | After visual stimulation (Miss licks) | Reconstructed H·W | Spearman + Wilcoxon signed-rank | 100 trials | < 2.22e-308 (rho); signrank p = 3.33e-02 *** | –0.227 | 0.630 |

#### Panel l — Cohen's d (Hit vs Miss, Expert) for Behavioural Variables in (k)

| Panel | Period | Variable | n recordings | Cohen's d [95% CI] | Sign-rank p | Above \ | d = 0.3? |
|---|---|---|---|---|---|---|---|
| l | Before Stimulus | Lick rate | 21 | +0.156 [+0.023, +0.297] | 0.0273 | No | Positive control; bounded |

| | | | | | | | |
|---|---|---|---|---|---|---|---|
| I | Before Stimulus | Pupil size | 21 | –0.155 [–0.341, +0.017] | 0.0853 | No | |
| I | Before Stimulus | Pupil dilation speed | 21 | –0.051 [–0.218, +0.104] | 0.476 | No | |
| I | Before Stimulus | Pupil position azimuth | 21 | +0.059 [–0.075, +0.196] | 0.566 | No | |
| I | Before Stimulus | Pupil position elevation | 21 | –0.032 [–0.170, +0.107] | 0.543 | No | |
| I | Before Stimulus | Locomotion | 4 | +0.190 [–0.365, +0.540] | 0.875 | No | Few recordings with sufficient locomotion variance |
| I | During Stimulus | Lick rate | 22 | **+2.774 [+2.524, +3.037]** | 4.01e-05 | **Yes** | Large effect — only behavioural variable above equivalence bound |
| I | During Stimulus | Pupil size | 21 | –0.235 [–0.396, –0.085] | 0.0106 | No | CI within equivalence band |
| I | During Stimulus | Pupil dilation speed | 21 | +0.010 [–0.145, +0.165] | 0.931 | No | |
| I | During Stimulus | Pupil position azimuth | 21 | +0.089 [–0.112, +0.288] | 0.394 | No | |
| I | During Stimulus | Pupil position elevation | 21 | –0.075 [–0.252, +0.103] | 0.394 | No | |
| I | During Stimulus | Locomotion | 4 | +0.101 [–0.403, +0.605] | 0.875 | No | Few recordings with sufficient locomotion variance |

### Extended Data Figure 8 — Binarisation Controls

#### Panel c — Dice Similarity Coefficient: Threshold 2.0 vs Alternative Thresholds

| Panel | Comparison | Test | n | p-value | Hedges' g (paired) |
|---|---|---|---|---|---|
| c | Across all 5 thresholds (omnibus) | Friedman | n = 758 trials × 5 thresholds | < 2.22e-308 *** | — |
| c | Threshold 2.0 vs 0.5 dF/F | Wilcoxon signed-rank (paired) | n = 758 trials | 2.59e-108 *** | +1.318 |
| c | Threshold 2.0 vs 1.0 dF/F | Wilcoxon signed-rank (paired) | n = 758 trials | 1.29e-29 *** | +0.424 |
| c | Threshold 2.0 vs 3.0 dF/F | Wilcoxon signed-rank (paired) | n = 758 trials | 8.82e-90 *** | +0.981 |
| c | Threshold 2.0 vs 5.0 dF/F | Wilcoxon signed-rank (paired) | n = 758 trials | 8.71e-111 *** | +1.512 |

### Extended Data Figure 9 — Parameter Search and Fits

#### Panel I — WD vs Radial Distance from Grid Centre (Spatial Homogeneity)

| Panel | Comparison | Test | n | p-value | Slope |
|---|---|---|---|---|---|
| I | Wasserstein distance to ground truth vs radial distance from grid centre | Pearson correlation + linear regression | n = 741 voxel positions (19 × 39 analysis-grid windows) | < 0.001 (annotated p = 0.000) *** | ≈ 0.0000 |

### Extended Data Figure 10 — Parameter Correlations

#### Panel a — Coupling strength c across conditions

Pairwise Wilcoxon rank-sum (Mann-Whitney U) on the 10 top-match parameter sets per condition; Kruskal-Wallis omnibus over all four conditions. Significance reflects the raw p-value.

| Panel | Comparison | Test | n | p-value | Hedges' g | Significance |
|---|---|---|---|---|---|---|
| a | c across Naive/Beginner/Expert/NoSpout | Kruskal-Wallis | 40 (10/cond) | 6.47e-01 n.s. | — | n.s. |
| a | c: Naive vs Beginner | Mann-Whitney U | 10 vs 10 | 8.10e-01 n.s. | –0.31 | n.s. |
| a | c: Naive vs Expert | Mann-Whitney U | 10 vs 10 | 2.31e-01 n.s. | –0.58 | n.s. |
| a | c: Naive vs NoSpout | Mann-Whitney U | 10 vs 10 | 2.62e-01 n.s. | –0.56 | n.s. |
| a | c: Beginner vs Expert | Mann-Whitney U | 10 vs 10 | 6.93e-01 n.s. | –0.19 | n.s. |
| a | c: Beginner vs NoSpout | Mann-Whitney U | 10 vs 10 | 6.93e-01 n.s. | –0.19 | n.s. |
| a | c: Expert vs NoSpout | Mann-Whitney U | 10 vs 10 | 1.00e+00 n.s. | +0.00 | n.s. |

#### Panel b — Decay constant d across conditions

| Panel | Comparison | Test | n | p-value | Hedges' g | Significance |
|---|---|---|---|---|---|---|
| b | d across Naive/Beginner/Expert/NoSpout | Kruskal-Wallis | 40 (10/cond) | 5.66e-01 n.s. | — | n.s. |
| b | d: Naive vs Beginner | Mann-Whitney U | 10 vs 10 | 3.39e-01 n.s. | +0.49 | n.s. |
| b | d: Naive vs Expert | Mann-Whitney U | 10 vs 10 | 1.00e+00 n.s. | +0.00 | n.s. |
| b | d: Naive vs NoSpout | Mann-Whitney U | 10 vs 10 | 8.47e-01 n.s. | –0.13 | n.s. |
| b | d: Beginner vs Expert | Mann-Whitney U | 10 vs 10 | 2.07e-01 n.s. | –0.54 | n.s. |
| b | d: Beginner vs NoSpout | Mann-Whitney U | 10 vs 10 | 3.01e-01 n.s. | –0.63 | n.s. |
| b | d: Expert vs NoSpout | Mann-Whitney U | 10 vs 10 | 6.18e-01 n.s. | –0.14 | n.s. |

##### Panel c — Inhibition range r across conditions

| Panel | Comparison | Test | n | p-value | Hedges' g | Significance |
|---|---|---|---|---|---|---|
| c | r across Naive/Beginner/Expert/NoSpout | Kruskal-Wallis | 40 (10/cond) | 8.76e-02 n.s. | — | n.s. |
| c | r: Naive vs Beginner | Mann-Whitney U | 10 vs 10 | 3.02e-02 * | –1.02 | * (raw); n.s. (Holm) |
| c | r: Naive vs Expert | Mann-Whitney U | 10 vs 10 | 2.62e-01 n.s. | –0.69 | n.s. |
| c | r: Naive vs NoSpout | Mann-Whitney U | 10 vs 10 | 9.34e-01 n.s. | +0.12 | n.s. |
| c | r: Beginner vs Expert | Mann-Whitney U | 10 vs 10 | 6.39e-01 n.s. | +0.15 | n.s. |
| c | r: Beginner vs NoSpout | Mann-Whitney U | 10 vs 10 | 2.13e-02 * | +1.19 | * (raw); n.s. (Holm) |
| c | r: Expert vs NoSpout | Mann-Whitney U | 10 vs 10 | 3.01e-01 n.s. | +0.80 | n.s. |

##### Panel d — Bias h across conditions

| Panel | Comparison | Test | n | p-value | Hedges' g | Significance |
|---|---|---|---|---|---|---|
| d | h across Naive/Beginner/Expert/NoSpout | Kruskal-Wallis | 40 (10/cond) | 2.81e-01 n.s. | — | n.s. |
| d | h: Naive vs Beginner | Mann-Whitney U | 10 vs 10 | 9.52e-02 n.s. | +0.71 | n.s. |
| d | h: Naive vs Expert | Mann-Whitney U | 10 vs 10 | 1.96e-01 n.s. | +0.54 | n.s. |
| d | h: Naive vs NoSpout | Mann-Whitney U | 10 vs 10 | 2.83e-01 n.s. | +0.34 | n.s. |
| d | h: Beginner vs Expert | Mann-Whitney U | 10 vs 10 | 5.54e-01 n.s. | –0.28 | n.s. |
| d | h: Beginner vs NoSpout | Mann-Whitney U | 10 vs 10 | 3.62e-01 n.s. | –0.44 | n.s. |
| d | h: Expert vs NoSpout | Mann-Whitney U | 10 vs 10 | 4.93e-01 n.s. | –0.21 | n.s. |

---

##### Panel e — Pairwise Pearson correlations within each condition (top-10 parameter sets, n = 10 per condition)

The 4 heat-maps in panel e display the 5 × 5 symmetric correlation matrix for each condition; the 10 unique off-diagonal pairs are reported below.

| Panel | Condition | Pair | n | r | p-value |
|---|---|---|---|---|---|
| e | Naive | β ↔ c | 10 | +0.093 | 7.98e-01 n.s. |
| e | Naive | β ↔ d | 10 | +0.199 | 5.82e-01 n.s. |
| e | Naive | β ↔ r | 10 | +0.527 | 1.17e-01 n.s. |
| e | Naive | β ↔ h | 10 | +0.147 | 6.85e-01 n.s. |
| e | Naive | c ↔ d | 10 | +0.762 | 1.04e-02 * |
| e | Naive | c ↔ r | 10 | +0.110 | 7.62e-01 n.s. |
| e | Naive | c ↔ h | 10 | +0.246 | 4.93e-01 n.s. |
| e | Naive | d ↔ r | 10 | –0.069 | 8.50e-01 n.s. |
| e | Naive | d ↔ h | 10 | +0.799 | 5.54e-03 ** |
| e | Naive | r ↔ h | 10 | –0.287 | 4.21e-01 n.s. |
| e | Beginner | β ↔ c | 10 | +0.670 | 3.39e-02 * |
| e | Beginner | β ↔ d | 10 | +0.203 | 5.73e-01 n.s. |
| e | Beginner | β ↔ r | 10 | –0.283 | 4.29e-01 n.s. |
| e | Beginner | β ↔ h | 10 | –0.460 | 1.81e-01 n.s. |
| e | Beginner | c ↔ d | 10 | +0.630 | 5.07e-02 n.s. |
| e | Beginner | c ↔ r | 10 | –0.144 | 6.90e-01 n.s. |
| e | Beginner | c ↔ h | 10 | –0.448 | 1.94e-01 n.s. |
| e | Beginner | d ↔ r | 10 | +0.455 | 1.86e-01 n.s. |
| e | Beginner | d ↔ h | 10 | +0.381 | 2.77e-01 n.s. |
| e | Beginner | r ↔ h | 10 | +0.606 | 6.36e-02 n.s. |
| e | Expert | β ↔ c | 10 | +0.309 | 3.85e-01 n.s. |
| e | Expert | β ↔ d | 10 | +0.396 | 2.57e-01 n.s. |
| e | Expert | β ↔ r | 10 | –0.188 | 6.04e-01 n.s. |
| e | Expert | β ↔ h | 10 | +0.302 | 3.97e-01 n.s. |
| e | Expert | c ↔ d | 10 | +0.791 | 6.39e-03 ** |

| | | | | | |
|---|---|---|---|---|---|
| e | Expert | c ↔ r | 10 | –0.613 | 5.95e-02 n.s. |
| e | Expert | c ↔ h | 10 | +0.100 | 7.84e-01 n.s. |
| e | Expert | d ↔ r | 10 | –0.825 | 3.34e-03 ** |
| e | Expert | d ↔ h | 10 | +0.679 | 3.10e-02 * |
| e | Expert | r ↔ h | 10 | –0.622 | 5.48e-02 n.s. |
| e | NoSpout | β ↔ c | 10 | –0.314 | 3.77e-01 n.s. |
| e | NoSpout | β ↔ d | 10 | –0.470 | 1.70e-01 n.s. |
| e | NoSpout | β ↔ r | 10 | +0.772 | 8.87e-03 ** |
| e | NoSpout | β ↔ h | 10 | –0.647 | 4.32e-02 * |
| e | NoSpout | c ↔ d | 10 | +0.900 | 3.87e-04 *** |
| e | NoSpout | c ↔ r | 10 | –0.339 | 3.38e-01 n.s. |
| e | NoSpout | c ↔ h | 10 | +0.457 | 1.84e-01 n.s. |
| e | NoSpout | d ↔ r | 10 | –0.508 | 1.34e-01 n.s. |
| e | NoSpout | d ↔ h | 10 | +0.785 | 7.16e-03 ** |
| e | NoSpout | r ↔ h | 10 | –0.610 | 6.10e-02 n.s. |

#### Panel f — Pooled pairwise Pearson correlations across all conditions (n = 40 = 10 × 4)

| Panel | Pair | n | r | p-value |
|---|---|---|---|---|
| f | β ↔ c | 40 | –0.047 | 7.72e-01 n.s. |
| f | β ↔ d | 40 | +0.010 | 9.52e-01 n.s. |
| f | β ↔ r | 40 | –0.202 | 2.11e-01 n.s. |
| f | β ↔ h | 40 | +0.052 | 7.51e-01 n.s. |
| f | c ↔ d | 40 | +0.723 | 1.39e-07 *** |
| f | c ↔ r | 40 | –0.226 | 1.60e-01 n.s. |
| f | c ↔ h | 40 | –0.007 | 9.65e-01 n.s. |
| f | d ↔ r | 40 | –0.320 | 4.41e-02 * |
| f | d ↔ h | 40 | +0.664 | 3.03e-06 *** |
| f | r ↔ h | 40 | –0.228 | 1.57e-01 n.s. |
